# Single neurons in thalamus and subthalamic nucleus process cardiac and respiratory signals in humans

**DOI:** 10.1101/2022.07.25.501133

**Authors:** Emanuela De Falco, Marco Solcà, Fosco Bernasconi, Mariana Babo-Rebelo, Nicole Young, Francesco Sammartino, Ali Rezai, Vibhor Krishna, Olaf Blanke

## Abstract

Visceral signals are constantly processed by our central nervous system, enable homeostatic regulation, and influence perception, emotion, and cognition. While visceral processes at cortical level have been extensively studied using non-invasive imaging techniques, very few studies have investigated how this information is processed at the single neuron level, both in humans and animals. Subcortical regions, relaying signals from peripheral interoceptors to cortical structures, are particularly understudied and how visceral information is processed in thalamic and subthalamic structures remains largely unknown. Here, we took advantage of intraoperative microelectrode recordings in patients undergoing surgery for deep brain stimulation (DBS) to investigate the activity of single neurons related to cardiac and respiratory functions in three subcortical regions: Ventral Intermedius nucleus (Vim) and Ventral caudalis nucleus (Vc) of the thalamus, and subthalamic nucleus (STN). We report that the activity of a large portion of the recorded neurons (about 70%) was modulated by either the heartbeat, the cardiac inter-beat interval, or the respiration. These cardiac and respiratory response patterns varied largely across neurons both in terms of timing and their kind of modulation. We observed neurons with increases or decreases in firing rate in response to either the heartbeat or the inter-beat interval. Peaks of neural activity were found at different phases of the cardiac and respiratory cycles. Whereas most neurons only responded to one of the tested signals, a substantial proportion of these visceral neurons (30%) was responsive to more than one of the tested signals, underlining specialization and integration of cardiac and respiratory signals in STN and thalamic neurons. By extensively describing for the first time single unit activity related to cardiorespiratory function in thalamic and subthalamic neurons, our results highlight the major role of these subcortical regions in the processing of visceral signals.

## Introduction

Our brain continuously receives signals originating from visceral organs providing a moment-by-moment sense of the physiological condition of the body. Such monitoring of bodily organs ensures the stability of the organism through regulation of homeostatic reflexes and adaptive behavior (Berntson et al., 1993; Damasio & Carvalho, 2013). There has been a recent upsurge of interest in the understanding of how information from internal organs is integrated in the human brain, because, beyond its function in homeostasis, visceral processing and the perception of the physiological condition of the body (i.e. interoception) are increasingly recognized to impact cognitive, social, and affective processes (Blanke et al., 2015; Critchley & Garfinkel, 2017; Critchley & Harrison, 2013; Engelen et al., 2023; Park et al., 2014; Seth, 2013; Tsakiris & Critchley, 2016) and to be a prominent feature of several psychiatric disorders (Khalsa et al., 2018; Khalsa & Lapidus, 2016; Murphy et al., 2017; Nord & Garfinkel, 2022).

Based on animal studies relying on electrical stimulation, chemical activation, pharmacological interventions, and various tracing methods it is commonly acknowledged that visceral inputs reach the central nervous system through the vagal nerve or the spinothalamic tract projecting to the nucleus tractus solitarius, the parabrachial nucleus, and periaqueductal gray matter (Berntson & Khalsa, 2021; Cechetto, 2014; Craig, 2009; Saper, 2002). Importantly, chest movement or pulsatility of blood vessels are also detected by proprioceptive or tactile peripheral receptors and are conveyed in the dorsal column-medial lemniscus pathway. Visceral information is subsequently transmitted to the hypothalamus, the amygdala, and the thalamus, from which it reaches primary sites of viscerosensory cortex such as the insula and the anterior cingulate cortex (Critchley & Harrison, 2013). Specifically, the ventral posterior complex of the thalamus has been put forward as the main relay of visceral information traveling to the cortex in most animal models (Cechetto & Saper, 1987; Groenewegen & Witter, 2004). Importantly, electrophysiological studies in animals have shown single unit responses to thermal (Emmers, 1966), gustatory (Emmers, 1966; Frommer, 1961), gastrointestinal (Asato & Yokota, 1989; Bruggemann et al., 1994), respiratory (Chen et al., 1992; Vibert et al., 1979), and cardiac (Massimini et al., 2000) activity located in ventral posterior medial (VPM) (Emmers, 1966; Frommer, 1961), ventral posterior lateral (VPL) (Asato & Yokota, 1989; Bruggemann et al., 1994), and/or posterior part of the Ventro Lateral Nucleus (VLp) of the thalamus. Similarly, STN neurons in the cat have been reported to tonically fire during specific respiratory phases (Vibert et al., 1979) and, when stimulated, to induce strong cardiorespiratory responses (Eldridge et al., 1981, 1985).

In humans, visceral processing has been characterized in different cortical and limbic regions, while the interoceptive ascending pathways linking the periphery to higher-order regions of the brain have been relatively neglected (Berntson & Khalsa, 2021). One extensively studied signal is the Heartbeat Evoked Potential (HEP) - obtained by averaging surface EEG signals time-locked to heartbeats - which has been associated with widely distributed cortical sources over frontal, central and parietal regions (see Park & Blanke, 2019a for a recent review). HEP has been linked with cardiac function (MacKinnon et al., 2013; Schandry & Montoya, 1996), but also with various cognitive and affective processes (Gray et al., 2007; Montoya et al., 1993; Park et al., 2014, 2016, 2018; Pollatos & Schandry, 2004; Schandry et al., 1986; Shao et al., 2011; Solcà et al., 2020). Although less investigated than cardiac signals, respiratory signals have also been shown to modulate brain oscillations at rest in widespread brain networks (Betka et al., 2022; Kluger & Gross, 2021). Similarly to cardiac signals, respiration has also been shown to influence a range of cognitive and motor processes (Adler et al., 2014; Allard et al., 2017; Kluger et al., 2021; Park et al., 2020; Perl et al., 2019; Rassler & Raabe, 2003; Schulz et al., 2016; Zelano et al., 2016).

A powerful approach to characterize how visceral information is encoded in the human brain is to use invasive recordings that directly measure local neural activity. Recently, such studies have profited from the high spatial and temporal resolution of intracranial electroencephalography (iEEG) and local field potentials to reveal specific heart-related processes in primary somatosensory cortex (Kern et al., 2013), but also insula, operculum, amygdala and fronto-temporal cortex (Park et al., 2018). Slow (∼0.2 Hz) respiration-entrained oscillatory activity has also been reported in the human limbic system (Herrero et al., 2018; Zelano et al., 2016). However, at the single neuron level, to date only two studies have been able to describe a relation between cardiac signals (i.e. duration of the cardiac cycle) and neural firing rates, reporting evidence for cardiac coding in human anterior cingulate cortex and medial temporal lobe (Frysinger & Harper, 1990; K. Kim et al., 2019). For respiration, while prior intracranial studies in humans have shown local neural activity linked to breathing (Dlouhy et al., 2015; Seyal & Bateman, 2009), to date no study has, to the best of our knowledge, investigated the link between respiration and single neuron activity in humans.

However, major brainstem and midbrain centers are involved in the processing and regulation of cardiac and respiratory signals for homeostasis and control of adaptive behavior (Berntson & Khalsa, 2021). While cortical processing of visceral signals has thus been elucidated by the listed studies, a fundamental gap in our understanding of visceral processing in the human brain remains. How are visceral signals processed and relayed at subcortical centers that project to the above-mentioned cortical structures? The neural properties of subcortical relays, such as the thalamic and subthalamic nuclei, are particularly understudied, likely due to the methodological challenges in accessing and analyzing these signals in humans. Rare human neuroimaging experiments (i.e. fMRI) and neurostimulation protocols have suggested an involvement of the mediodorsal thalamus, the periaqueductal gray matter, the STN, the hypothalamus in autonomic system regulation (Beissner et al., 2013; Thornton et al., 2002), and of different subregions of the thalamus in respiratory control (Faull et al., 2015; K. Pattinson et al., 2009). Moreover, electrical stimulation of the ventral portion of the human thalamus (Thornton et al., 2002) and STN (Sverrisdóttir et al., 2014) modulates baroreceptors sensitivity, increases heart rate and blood pressure while changes in single neuron activity related to cardio-vascular modulation have been reported in the ventro-postero-lateral nucleus of the thalamus (Oppenheimer et al., 1998) and STN (Coenen et al., 2008).

An invaluable opportunity to investigate the subcortical functional aspects of visceral responses in humans are intraoperative recordings of single neurons during the implantation procedure of electrodes for deep brain stimulation (DBS). During DBS surgeries, intraoperative microelectrode recordings (MER) provide the opportunity to record from thalamic and subthalamic nuclei in awake humans (Marks, 2015). However, an important challenge in DBS recordings - especially when investigating signals time-locked to the heart - is the presence of pulsatility artifacts (PA) resulting from the pulsating motion of the brain and nearby pulsating blood vessels. Such movements lead to changes in the recorded spike waveform across the cardiac cycle that can affect spike detection and bias neuron classification. To our knowledge, only one study, so far, investigated single cell activity related to cardio-vascular processing in the human thalamus (Oppenheimer et al., 1998). While representing an important and pioneering report, Oppenheimer and colleagues limited their investigation to the relationship of thalamic neuronal activity with respect to the pulse signal (measured via a pulse-oximeter on the toe; as no ECG was available for the analyzed dataset). Furthermore, single units and their stability were only assessed by visual inspection, which may lead to multi-unit clusters and result in an underestimation of the number of modulated units.

Here, we combine MER with simultaneous physiological signal acquisition (heart rate, HRV, breathing rate, breathing variability) to investigate whether single neurons in the thalamus and STN process basic signals of heartbeat and respiration. To overcome the challenges of intraoperative MER recordings faced by Oppenheimer and colleagues, we applied latest single unit analysis methodology, using unsupervised parametric spike sorting algorithms (Chaure et al., 2018; Quian Quiroga et al., 2004) and a recently developed method for the analysis of cardiac-motion features in the recoded spikes (Mosher et al., 2020). Following preprocessing of the extracellular potentials, we identified three different visceral signals in thalamic and subthalamic neurons, linked to (1) the heartbeat, (2) the duration of the cardiac cycle, and (3) the respiratory cycle. The activity of a large portion of the recorded neurons (about 70%) was modulated by at least one of these visceral signals, with a substantial proportion of those neurons (30%) encoding multiple signals. Our results provide the first functional characterization of the activity of thalamic and subthalamic single neuron populations in humans encoding cardiorespiratory activity, highlighting the major role of these subcortical regions in interoceptive processing.

## Results

### Microelectrodes recording and neurons identification

We recorded data during DBS implantation surgery in a total of 23 patients. Implantation site of all recordings was based solely on medical indication, that is Vim target for essential tremor and STN target for Parkinson disease. We acquired data from three different subcortical regions: the subthalamic nucleus (STN), the ventral intermedius (Vim) thalamic nucleus, and ventral caudal (Vc) thalamic nucleus. Figure 1 shows the MNI normalized coordinates of the recording location across the 23 surgeries (63 recording sessions in total).

**Figure 1.**
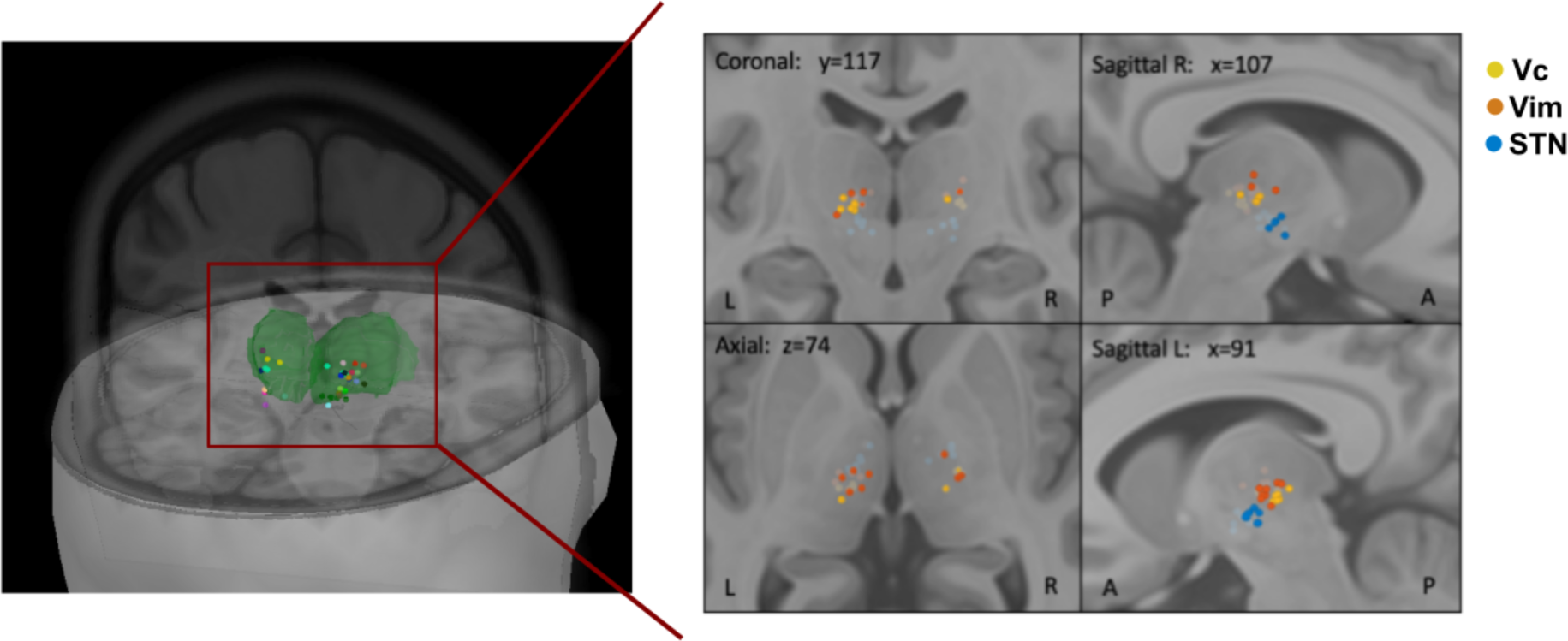
Visceral processing pathways and recordings. Location of recording electrodes across the 23 surgeries color-coded by patient (left) and color-coded by identified region for Vc (yellow), Vim (orange), STN (blue) (right). Green volume represents the thalamus. Labeling of recording sites was obtained intraoperatively based on each patient’s brain scan, coordinates were then converted to MNI space for visualization and are projected here on the MNI/ICBM152 human brain template (Mazziotta et al., 1995). Note that for some sessions, location dots overlap as different recordings were performed in the same location. Abbreviations used: ventral caudal thalamic nucleus (**Vc**), ventral intermedius thalamic nucleus (**Vim**), subthalamic nucleus (**STN**). Left and right panels generated using Brainstorm (Tadel et al., 2011).

After the initial spike detection and sorting we identified 127 putative single units, of which 27 were in STN, 45 in Vc, and 55 in Vim (mean number of units per recording session = 2.0, SD: 1.1). We then applied a modified version of a recently proposed method (Mosher et al., 2020) to identify and remove neurons that are contaminated by cardiac-related movement artifacts in their EAPs (see Methods and supplementary Figure S1). Following this procedure, 73 neurons were considered for further analysis (16 in STN, 24 in Vc, and 33 in Vim). The number of neurons detected in each area per patient are reported in Table S1.

### Neural responses to heartbeat

Modulation of the neural activity in response to the heartbeat was assessed by analyzing the changes in neural firing rates along the cardiac cycle for each of the 73 neurons (following exclusion of neurons with artifacts) by computing a non-uniformity test and the coherence between spikes and cardiac phase (see Methods for further details). This analysis led to the identification of 30 neurons (41%) with a significant modulation of their firing activity along the cardiac cycle. In a post-hoc analysis, we tested whether the proportion of responding neurons differed between recordings sites and found a significantly larger proportion of neurons responded to the heartbeat in both STN (10 units, 63%) and Vim (16 units, 48%) compared to Vc (4 units, 17%) (z-test on proportions, STN vs Vc: *z*(38) = 3.1, *p* = 0.002; Vim vs Vc: *z*(55) = 2.7, *p* = 0.006). Some exemplary responsive units across STN, Vim, and Vc are shown in Figure 2A. Neurons presented different response patterns along the cardiac cycle. For example, the STN neuron in Figure 2A (left panel) shows a clear decrease in firing activity following the R-peak, with the average number of spikes approaching zero during the central part of the cardiac cycle. Conversely, the illustrative Vc and Vim neurons shown in the central and right panel of Figure 2A show a different pattern, that is a marked increase in spiking activity following the R-peak. Across the 30 responsive units, 13 increased their firing rate following the R-peak (43%) and 17 decreased it (57%). No significant difference in the proportion of increasing vs decreasing responses was observed (z-test, *z* (28) = 1.02, *p* = 0.3). In addition, when we repeated our analysis by time-locking the spikes to the end of the cardiac cycle (i.e., looking backwards from the R-peak), we found that all the 30 neurons still exhibited a significant response. Furthermore, we verified that the identified heartbeat responses could not be explained by spikes near the detection threshold (supplementary results, Figure S5).

**Figure 2.**
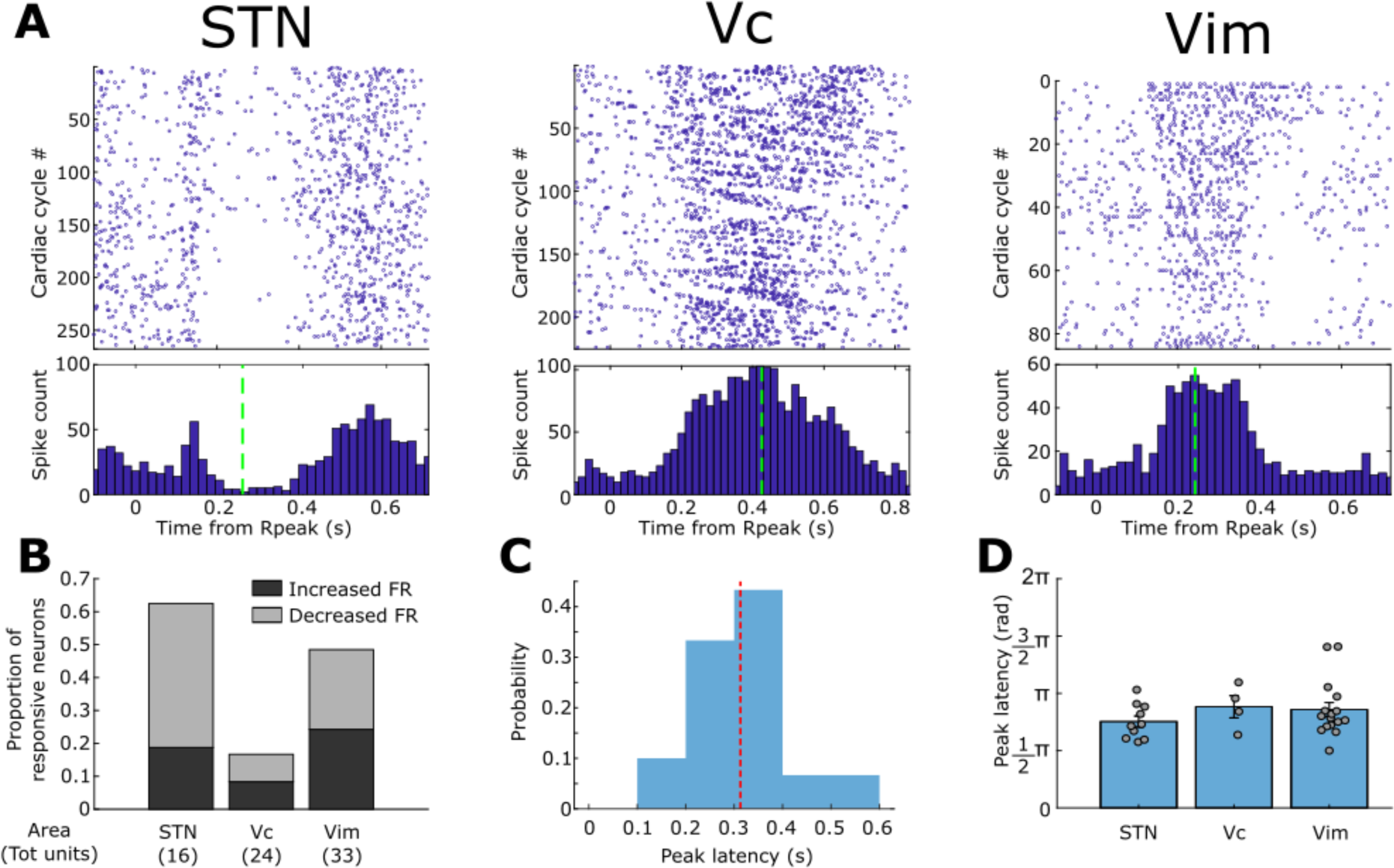
Neural responses to heartbeats. **A)** Examples of neurons responding to the heartbeat with firing rate modulations in the three different target regions (STN, Vc and Vim) recorded from three different patients. Top panel shows the raster plot for the neural spikes aligned to the R-peak across the different cardiac cycles. Bottom plot shows the cumulative histogram of spikes across cardiac cycles binned in 10ms bins. Green dotted lines represent the peak latency (time with maximal modulation of the response). Maximum value for the x-axis is equal to the median IBI for the recording session. Sample recordings and EAP waveform modulation test for these exemplary units are shown in Figures S4.1, S4.2, and S4.3 **B)** Proportion of neurons showing a significant increase (dark gray) or decrease (light gray) of firing rates along the cardiac cycle in the three regions. The total number of neurons tested in each region is shown in parentheses. **C)** Distribution of peak latencies for the 30 responsive neurons identified. Average latency was 313ms (SD=100ms). 77% of responses had a peak latency between 200 and 400ms. **D**) Comparison of peak latencies normalized by IBI (in rad) across areas. Bars represent the mean (±SEM). Circles represent single peak latencies measured. No significant difference in peak latency was observed across regions (ANOVA test, *F*(2, 27) = 0.89, *p* = 0.42). Figure 2B shows the proportion of significant responsive neurons identified in the three regions color-coded per type of response (increased or decreased firing). Although, the STN was found to present the largest proportion of decreasing responses (70%), compared to the two thalamic nuclei (50% both in Vc and Vim), the differences in proportion were not significant (z-test on STN vs thalamus (Vc + Vim), *z* (28) = 1.04, *p* = 0.29).

For each response, the peak latency (represented by the green dotted lines in Figure 2A) from the ECG’s R-peak was determined from the instantaneous firing rates as the time where the modulation of activity (increase or decrease) was maximal. Distribution of all peak latencies for the 30 responsive neurons is shown in Figure 2C. The average peak latency across responses was 313ms (SD = 100ms), with 77% of peak latencies (23 out of 30) falling in the interval between 200 and 400ms. In terms of angular latency (in rad), the mean latency was 2.5rad (SD = 0.6rad). Peak latencies across regions were compared in terms of their angular values (in rad) that corrects for differences in IBI across recordings. No significant difference was observed across regions according to an ANOVA test (*F*(2, 27) = 0.89, *p* = 0.42) (Figure 2D). Furthermore, no significant difference in peak latencies was observed between increasing and decreasing responses (unpaired t-tests: *t*(28) = 0.53, *p* = 0.6).

These data show that signals about the cardiac cycle are widely encoded in thalamic and subthalamic neurons (41% of units). These signals manifest as a rhythmic modulation (increase/decrease) of spiking activity during specific portions of the cycle, with the peak modulation on average around 300ms (post R-Peak), roughly corresponding to the end of the first half of the cardiac cycle.

### Relationship between firing rates and Inter-beat Interval (IBI)

The heart rate variability (HRV) – the fluctuation in the time intervals between consecutive heartbeats – is an important physiological marker, in cardiac physiology, of heart-brain interactions, and a useful tool to assess sympathetic and parasympathetic influences on disease states (Khan et al., 2019). We therefore asked whether human thalamic and sub-thalamic neurons process IBI-related signals. For each neuron, we tested the relationship between spiking activity and IBI by means of a Pearson correlation between the average firing rates during each cardiac cycle and the corresponding cycle duration (IBI). Figure 3A1 shows examples of neurons whose firing rate was significantly correlated with the IBI. The corresponding raster plots for the units are shown in Figure 3A2. Overall, we found significant correlation for 29% of the recorded units (21 out of 73, t-statistic corrected for multiple comparisons), with 9 units exhibiting a positive correlation (higher firing rates for longer IBIs) and 12 units exhibiting a negative correlation (lower firing rates for longer IBIs) (Figure 3B). The probability of positively/negatively correlated units across the three recorded regions is shown in Figure 3C. These data show that spontaneous spiking activity in the recorded regions is linked to IBI for almost one third of recorded neurons, and that longer IBIs can be linked to either positive or negative changes in firing rates.

**Figure 3.**
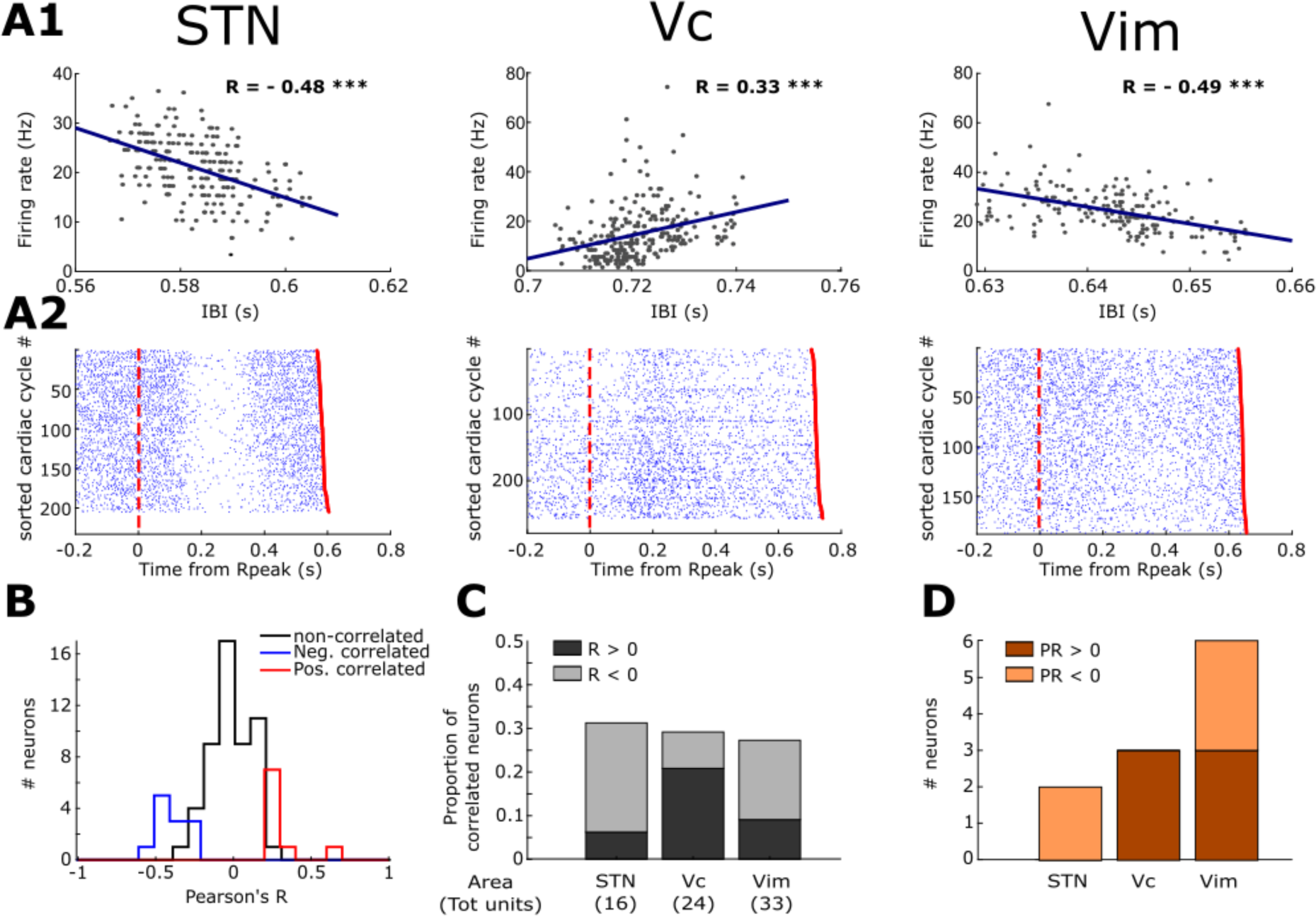
Neural activity covaries with Inter-Beat Interval. **A1)** Correlation between the duration of the cardiac cycle (IBI) and the firing rate across cycles in three exemplary neurons recorded from the three different target regions (STN, Vc and Vim). Pearson’s R is reported for each plot. Asterisks mark correlation significance (*** indicate *p*<0.001) **A2)** Raster plots from the three exemplary neurons of the neural spikes aligned to the R-peak across the different cardiac cycles where cycles were sorted according to their IBI. Red dotted line represents the beginning of the cardiac cycle (R-peak) while the red dot at the end of each trial represents the following R peak. **B)** Distributions of Pearson’s R values for the correlation between IBI and firing rates separated in un-correlated (black), and significantly correlated (blue for R<0, red for R>0). **C)** Proportion of recorded neurons exhibiting a significant positive (dark gray) or negative (light gray) correlation between IBI and firing rate in the three regions. The total number of neurons tested in each region is shown in parentheses. **D**) Number of recorded neurons in the three regions for which a significant phase-response (PR) of IBI to neural activity was detected. Bars are colored according to the sign of the detected PR (dark orange for positive, light orange for negative).

Given that the heart-brain communication is bidirectional and can be mediated by several pathways, we next tried to characterize the directionality of the observed relationship. We used phase-response (PR) analysis at the cycle by cycle level, as recently proposed by Kim and colleagues (K. Kim et al., 2019), to identify neurons for which the spiking activity elicited changes in the duration of the cardiac cycle. PR measures how much the IBI is shortened or lengthened in response to a neuron’s spiking activity. Among the units exhibiting a significant correlation between firing rate and IBI (21), we found that 11 also showed a significant PR of IBI to the spiking activity. Specifically, 6 units had positive PR (lengthening of cardiac cycle following spikes) and 5 units had a negative PR (shortening of cardiac cycle following spikes). As expected, the sign of PR was always matching the sign of Pearson’s R measured from the correlation.

Altogether, these results demonstrate that information about the duration of the cardiac cycle is encoded in the activity of about a third of the present thalamic and subthalamic neurons. Moreover, the relationship between firing rate and IBI was characterized either by a positive or negative correlation in STN, Vim, and Vc. In a few neurons, this relationship extended to more than one cardiac cycle (supplementary results, Figure S6).

### Neural activity is modulated by respiratory phase

Apart from the widely studied heart-related processes, interoception also embraces other systems, including input from the respiratory system. Respiratory-associated thalamic activity has been reported in cats (Chen et al., 1992), and the existence of respiratory signals in the human thalamus and STN would provide evidence for their broader implication in interoceptive processing. Furthermore, since respiratory signals have a much slower frequency than cardiac ones, their neural correlates are less prone to be affected by pulse artifacts. Therefore, we decided to capitalize on respiratory signals by analyzing the recorded neural activity in relationship to the respiratory cycle. Respiratory signals were extracted from the ECG traces using an ECG-Derived Respiration (EDR) algorithm (see Methods and Figure S7-A). We could successfully extract the EDR for all the recording session containing the identified neurons (see Methods). Given that systematic coupling between cardiac and respiratory signals exists (e.g. respiratory sinus arrhythmia), neurons showing a significant EAP modulation along the cardiac cycle were also excluded from all respiration-related analyses. Furthermore, we verified that none of the considered neurons had a significant EAP waveform modulation along the respiratory cycle either (see Methods). Therefore, all 73 neurons were considered for the following analysis. For each neuron, we assessed whether the spiking probability was non-uniform along the respiratory cycle, that is, if the neuron fired preferentially during a specific respiratory phase (preferred phase). The existence of a preferred phase was assessed via surrogate population testing that intrinsically corrects for eventual differences in the distribution of phases along the recording. Some examples of neurons that fired preferentially for a specific phase of the respiratory cycle are shown in Figure 4A1-A2. Overall, we identified 22 neurons (31%) whose activity was modulated by the respiratory phase (7 in STN, 5 in Vc, and 10 in Vim). The proportion of respiration-modulated neurons in each region is shown in Figure 4B. The distribution of preferred phases for those neurons is shown in Figure 4C. While we found that a slightly larger number of neurons (13 out of 22) fired preferentially during the inspiration (*π < φ < 2π*), no significant phase preference overall was identified (Rayleigh test for non-uniformity, *z*(21) = 0.37, *p*= 0.69). Additionally, the relationship between firing rates and duration of the respiratory cycle was tested similarly to what was done for the IBI (previous section). To this end we tested the correlation between average firing rate and the time interval from the beginning of one expiration to the beginning of the following one. None of the recorded units showed a significant correlation between the two measures.

**Figure 4.**
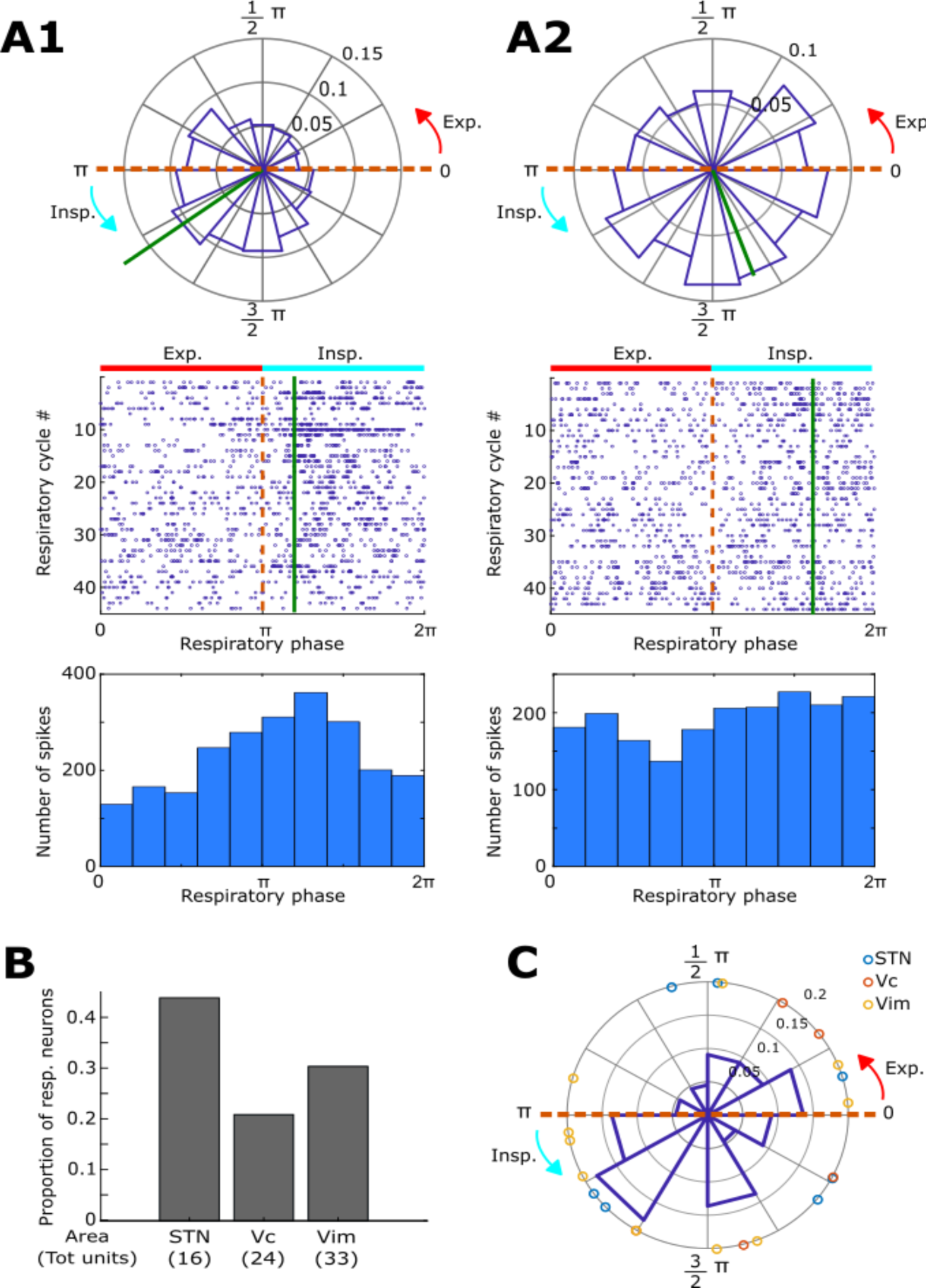
Neural activity is modulated by respiration. **A1)** Exemplary neuron in the STN whose spiking activity was significantly non-uniform along the respiratory cycle (surrogate test on Rayleigh statistic for non-uniformity, *p* = 10^-^4). Top: Circular histogram of spike density across the respiratory phase. The green line represents the mean vector of the distribution, with its angular direction being the preferred spiking phase for the neuron (φ = 3.8 rad). Middle: Raster plot for the neural spikes as a function of the respiratory phase across the different respiratory cycles. Bottom: cumulative histogram of spikes across respiratory cycles binned in π/5 rad bins. **A2)** Exemplary neuron in the Vim whose spiking activity was significantly non-uniform along the respiratory cycle (surrogate test on Rayleigh statistic for non-uniformity, *p* = 2*x*10^-3^). Conventions are as in panel B1. Preferred spiking phase φ = 5.1 rad **B)** Proportion of neurons whose activity was significantly modulated by the respiratory phase in the three recorded regions. The total number of neurons tested in each region is shown in parentheses. **C)** Distribution of preferred spiking phases for the 22 significantly modulated neurons. Circles indicate individual values (color-coded by region). No significant phase preference overall was identified (Rayleigh test for non-uniformity, *z*(21) = 0.37, *p*= 0.69).

These results show that information about the respiratory cycle is encoded in about one third of the thalamic and subthalamic neurons. While cortical neural excitability is known to vary over the respiratory cycle (Dulla et al., 2005), our findings cannot be simply explained by such a generalized increase of neural activity during a specific respiratory phase, because different neurons that we studied here exhibit different preferred phases for firing. Instead, the variability of preferred phases is more likely connected to the activity of respiratory neurons in the brainstem whose activation is observed in different phases of either inspiration or expiration (Onimaru et al., 1997).

### Overlap of interoceptive modalities

Given that we identified three different interoceptive signals in our dataset (heartbeat, cardiac IBI, respiration), we next asked whether the same responsive neurons encode these different types of signals or whether there are rather separate populations of neurons encoding each type of signal. When looking for neurons responding to each of the different signal types (Figure 5A) we found that most of them (35 out of 53 responsive units, 66%) responded to only a single type of interoceptive signal. From the remaining 18 neurons, 16 (30% of the total responsive units) responded to two types of stimuli, whereas only very few (2, 4%) responded to heartbeat, IBI, and respiration cycle. For example, the 21 neurons responding to the IBI were in the large majority distinct from those responding to the heartbeat (with only 7 exhibiting both heartbeat and IBI modulation), and distinct from those responding to respiration (with only 6 exhibiting both heartbeat and respiration modulation). The proportion of neurons that responded to each modality or to multiple modalities (multi-responsive visceral neurons) across the three recorded regions is shown in Figure 5B. There was no clear bias in the distribution of multi-responsive visceral neurons across regions. Furthermore, no relationship was found between respiratory sinus arrythmia, reflecting coupling between respiratory and cardiac signals, and multimodality of the responses (supplementary results, Figure S8).

**Figure 5.**
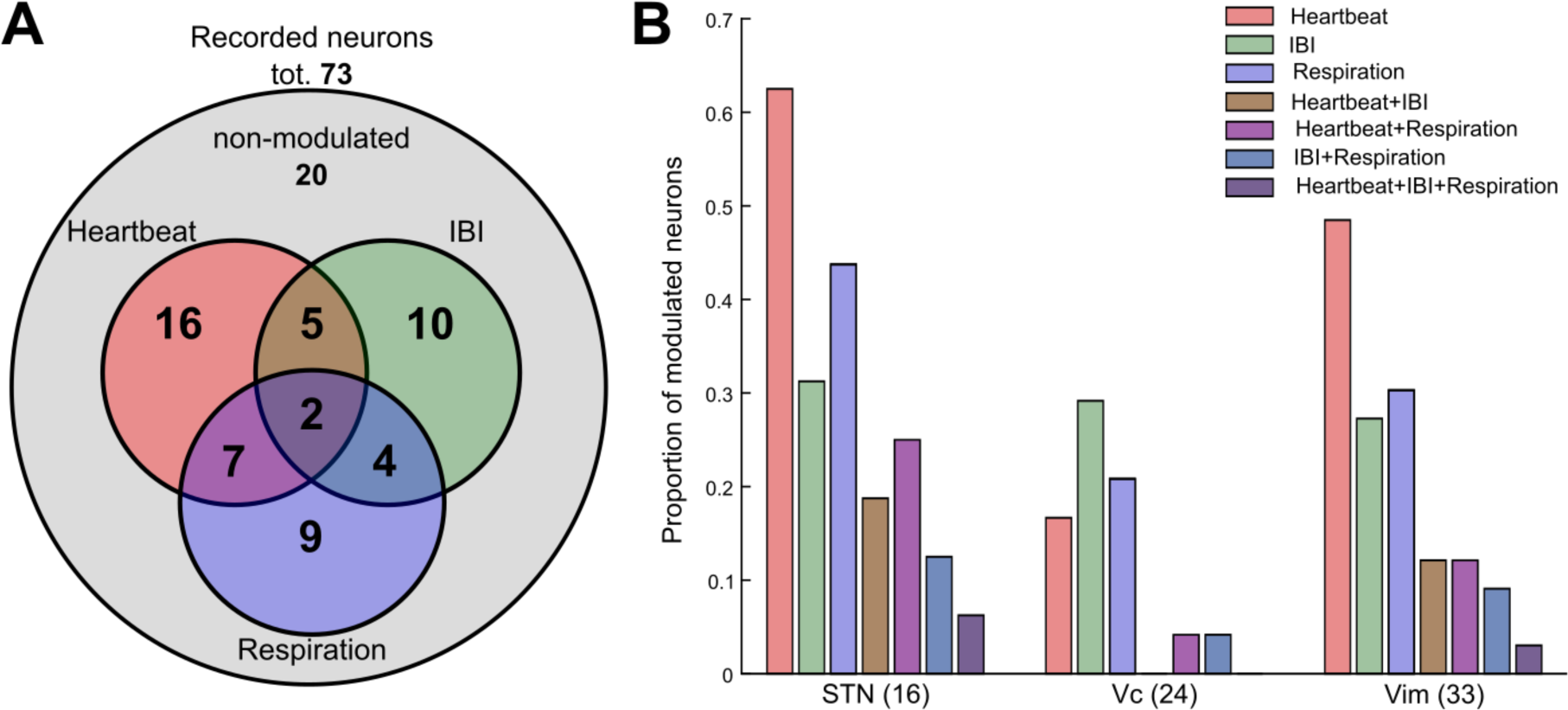
Numbers of different interoceptive neurons. **A)** Venn diagram showing the number of neurons among the recorded ones that responded to one or more interoceptive signals. Red circle: units responsive to the heartbeat; Green circle: units with firing rate correlated with IBI; Blue circle: units responsive to the respiratory phase. **B)** Proportion of different interoceptive neurons in each of the three regions color-coded by type of response. The total number of neurons tested in each region is shown in parentheses.

Overall, these data show that visceral signals are processed in the thalamus and STN by partially overlapping neural populations. The finding that diverse signals can be encoded by the same neuron underlies the interaction between different visceral modalities. Finally, these results are further reassurance that these cardiac and respiratory signals in thalamic and subthalamic neurons are not the effect of recording artifacts since the observed signals are independent from one another and not the result of the same underlying input.s

## Discussion

By acquiring intraoperative MER with simultaneous monitoring of physiological signals, we report that the activity of single neurons in thalamus and STN encodes visceral signals related to cardiac and respiratory function. In particular, we find that 53 out of the 73 single neurons included in the present analysis respond to at least one of the three interoceptive signals that we analyzed, that is around 40% responded to heartbeat, 30% to IBI, and 30% to the respiratory cycle. While a large proportion of these neurons were significantly modulated by cardiorespiratory activity, their patterns show variable response properties. We observed increases and decreases in firing rate, peaks of modulation during different phases of the respiratory and cardiac cycle, as well as positive or negative correlations between firing rate and the duration of the cardiac cycle. Finally, while most of the present neurons responded to a single type of signal, some units were found to encode two, or even all three tested interoceptive signals.

The present data significantly extend earlier preliminary evidence of neural responses to visceral inputs in the thalamus in human and non-human primates (Bruggemann et al., 1994; Chandler et al., 1992; Oppenheimer et al., 1998) by describing extensive neural responses to heartbeats, IBI, and respiratory cycle in the human thalamus. The thalamus is a major diencephalic structure bidirectionally relaying information between brainstems and cortical regions. Apart from olfaction, every sensory system transits to a thalamic nucleus that receives, processes, and sends information to a specific cortical area. For instance, the ventral posterolateral nucleus (equivalent to Vc according to the nomenclature of Hirai and Jones (Hirai & Jones, 1989)) receives medial lemniscal fibers from cuneate and gracile nuclei conveying somatosensory information to the somatosensory cortices. Similarly, visual inputs from the retina transit to the lateral geniculate nucleus before being further processed in the visual cortex. Thalamic neurons respond to a variety of stimuli, including tactile stimuli (skin mechanoreceptors) in the Vc (Lenz et al., 1993; Weiss et al., 2009) and retinal inputs in the dorsal lateral geniculate nucleus (Usrey & Alitto, 2015), and are driven by proprioceptive information (passive and active joint movement) in the Vim (Ohye et al., 1989; Raeva, 1986).

In addition to the thalamic nuclei Vim and Vc, we also recorded single cells in the STN – the most ventral part of the diencephalon situated between the thalamus and the midbrain which receives inputs from motor areas of the cerebral cortex, projects to the substantia nigra, and is reciprocally connected with the globus pallidus (Haines & Mihailoff, 2018). STN is classically considered as a motor structure given its important position in the basal ganglia circuitry and its prominent function in the execution and control of voluntary movements. Accordingly, neurons in the human STN have mainly been reported to be activated during different phases of sensorimotor tasks such as action selection, movement, and in response to stop signals (Mosher et al., 2021; Zavala et al., 2015). Our description of neuronal responses in STN to cardiac and respiratory inputs is thus at first sight surprising. However, the basal ganglia circuitry is also known to be involved in the control of autonomic function during respiration (independantly of voluntary breathing; Pazo & Belforte, 2002), and single neurons in the STN of cats have been reported to tonically fire in specific respiratory phases (Vibert et al., 1979). There is also evidence that STN is involved in cardiorespiratory control (Green & Paterson, 2020). Thus, midbrain stimulation (locomotor region in decorticated cats, corresponding to STN in humans)– induces strong cardiorespiratory responses (Eldridge et al., 1981, 1985). Moreover, changes in heart rate and blood pressure modulate spike activity in human STN (Coenen et al., 2008) and STN-DBS modulates baroreceptors sensitivity (a key regulator of cardiovascular homeostasis) (Sverrisdóttir et al., 2014) heart rate and blood pressure (Thornton et al., 2002). Extending this former evidence, here we report for the first time neurons in the human STN responding specifically to heartbeats, IBI and respiratory cycle. The neural responses observed in STN were similar to those observed in thalamus, both in terms of proportion of units modulated and in terms of variability of response patterns.

### Neural responses to heartbeats

In the present work, we found that the heartbeat is widely encoded in the human thalamus and the STN, that is about 40% of the neurons are significantly modulated by the cardiac cycle, with larger proportions in STN and Vim, compared to Vc (only 17% of Vc units exhibited modulation). The large proportion of neurons responding to the cardiac cycle seems reasonable given the significant proportions of cardiac-related activity observed in the hippocampus and amygdala of epileptic patients (∼20%) (Frysinger & Harper, 1990) and of other visceral information encoded in monkey thalamic cells (85%) (Bruggemann et al., 1994). However, a previous study in humans (Oppenheimer et al., 1998) only reported a very small percentage of thalamic units (<1%) exhibiting phasic modulation of firing rates related to cardio-vascular activity. While representing an important and pioneering report, this study had several important technical limitations. Firstly, neuronal spikes were only assessed by visual inspection; lack of spike sorting could have led to an underestimation of the single units recorded (because of multi-unit detection) and therefore of the units modulated. In addition, no ECG was recorded, and the authors only considered as responsive those neurons that showed a peak of activity aligned to the systolic peak pressure (measured via a pulse-oximeter on the toe) and at least 50% larger than the background firing rate. This differs from the present study where we used feature-based spike sorting and analyzed cyclical modulations (i.e. increase and decrease of firing rate) of neural activity along the entire cardiac cycle without setting an a-priori threshold for response, potentially explaining the present proportion of cardiac units. Furthermore, Oppenheimer et al. performed their recordings only in Vc, which showed the lowest number of responses to the heartbeat in our dataset (17%) with only two units showing an increase in firing rate following the R-peak. Finally, Oppenheimer et al. did not perform spike clustering and only 1 unit would have been detected in our dataset without this important step, which is consistent with this previous work.

The cardiac neurons we observed responded to the heartbeat by either decreasing (∼60% of responding units) or increasing (∼40% of responding units) their firing rate following the R peak. Decreases in neural firing rate (suppressive response) in response to external stimuli has been widely observed in other brain regions, such as the primary visual cortex and the inferior temporal cortex (Salehi et al., 2020). Decreasing responses in visual cortices for non-preferred stimuli are thought to be a mechanism that enhances tuning selectivity (Ringach et al., 2002). Accordingly, we propose that the increase/decrease of firing rate following R-peak may represent tuning to different periods of the cardiac cycle and that, likely, the entire cardiac cycle is tracked by thalamic and subthalamic neuronal populations. Alternatively, it is known that a multitude of systems and feedback loops exist between the different visceral brainstem and midbrain nodes and such inputs may also explain the simultaneous presence of both increasing and decreasing firing rate. We note that any conclusions about the directionality of the observed responses is difficult, considering that firing rate modulations were found when time-locking neurons spikes both to the R-peak and to the end of the cardiac cycle. This is likely the consequence of the extremely small variability of the IBI intervals in our recordings (Figure S2), possibly due monitored cardio-respiratory function (anesthesia during surgery) or to the movement disorder itself (Heimrich et al., 2021).

Concerning timing of activity, the peak of modulation for the neural activity for 77% of all units processing the heartbeat was found to fall between 200ms and 400ms post R-Peak, with a significant heterogeneity of latencies (SD = 100ms). This is consistent with animal data of subcortical visceral processing (Hayward & Felder, 1995). This timing suggests that for many of the identified neurons the modulation could potentially be linked to baroreceptor-related activations (given that the timing of blood ejection corresponds to ∼300ms after R-peak). This time window is also in line with the latency of HEP modulations (Kern et al., 2013) observed with both iEEG in insular recordings (Park et al., 2018) and with scalp EEG (MacKinnon et al., 2013). However, the significant variability of observed latencies indicates that the effects we observed are not limited to baroreceptor-related activation.

The large number of modulated neurons and their variable patterns of response underscore the importance and the complexity of the interplay between cardiac signals and brain function. Several processes and different information pass through STN and thalamus and could underlie the observed neural responses to heartbeats since different kinds of inputs reach the brain at each heartbeat. These different inputs include discharges from baroreceptors in the heart wall and blood vessels as well as signals from mechanoreceptors in the chest wall carrying information about the heart’s contraction timing and strength (Armour & Ardell, 2004). Additional signals include proprioceptive or tactile peripheral receptors which detect the pulsatility of blood vessels (Birznieks et al., 2012; Ford & Kirkwood, 2018; Macefield, 2003). Moreover, evidence also exists for astrocyte monitoring of cerebral perfusion and vasculo-neuronal coupling directly in the CNS (K. J. Kim et al., 2016; Marina et al., 2020). However, even higher levels of available oxygen could have increased the likelihood of spontaneous firing in the recorded neurons during the systole. Considering that we observed both increases and decreases in firing rates in different timing and given the sharpness of change in firing rate observed in many neurons, we argue that the multiple modulations we observed are most likely due to different mechanisms.

To summarize, we show that different phases of the cardiac cycle are encoded in a large proportion of thalamus and STN neurons. The timing of many neural responses observed is compatible with baroreceptor activation but the variability in latency and direction of the modulations indicates that multiple heartbeat-related signals are relayed in these subcortical structures.

### IBI-related signals

The IBI and its variability are important physiological parameters, as beat-to-beat monitoring and control of heart rate are fundamental to regulate cardiac function and ensure homeostasis in response to internal and external environmental changes (Khan et al., 2019). This precise regulation is maintained through local reflex and feedback loops involving afferent information related to cardiac function, cardiac frequency and blood pressure from various peripherical receptors (Bishop et al., 2011) that is relayed to subcortical and cortical structures. Descending projections to parasympathetic and sympathetic relays, in turn, regulate heart rate (chronotropic control), conduction (dromotropic control) and myocardial contractility (ionotropic control) (Berntson et al., 2007; Mccraty & Shaffer, 2015). However, very little is known about how single neuron activity in humans relates to the IBI. Two single cell studies in humans investigated this relationship at the cortical level and have reported a significant correlation between IBIs and the firing rates of 10% of anterior cingulate cortex neurons, and 20% of medial temporal lobe neurons (Frysinger & Harper, 1990; K. Kim et al., 2019). In the present study, when investigating the relationship between neural activity and the duration of the cardiac cycle, we found that the IBI is encoded in the activity of a substantial portion of thalamic and subthalamic neurons (about 30%), by showing that the firing rates of these neurons correlated with the IBI length. This is in line with evidence from animal research where the activity of 33% of somatosensory neurons in the cat thalamus was found to correlate with the IBI (Massimini et al., 2000). To our knowledge, the present data are the first report of such evidence in the human thalamus and STN, and show that all three tested regions are involved in neural IBI monitoring.

Importantly, we observed both positive (∼40%) and negative (∼60%) correlations, meaning that during longer cardiac cycles some neurons increased their firing rate, while others decreased it. Considering the bidirectionality and the complexity of homeostatic IBI control, these different responses most likely reflect different regulatory mechanisms. For instance, they could reflect the monitoring and processing of various signals (heart rate, cardiac function, blood pressure) to control the heart rate. Conversely, heart rate information not only influences chronotropic feedback control but also ionotropic and dromotropic parameters. We also note that the presence of both positive and negative correlations between firing rates and IBI rules out the trivial hypothesis that the different levels of oxygenation associated with the cardiac cycle duration might underlie the IBI effects (higher levels of oxygen available during shorter cardiac cycle, would only explain a generalized increase of neural activity, a negative correlation between firing rate and IBI). When investigating the directionality of the observed modulation using phase response analysis, we found that only in 50% of the IBI-correlated units the firing rate had a significant effect on the duration of the cardiac cycle (lengthening or shortening), suggesting that we observed an efferent signal in only half of the units. Using the same approach for neurons in the medial temporal lobe and anterior cingulate cortex, Kim and colleagues (K. Kim et al., 2019) reported a significant phase response for all units that exhibited a significant IBI-correlation. However, none of the units they identified showed a significant modulation of activity in response to the heartbeat itself, while we found that 7 out of 21 correlated units responded to the cardiac cycle.

To summarize, we observed an important number of neurons with firing rates that positively or negatively correlated with IBI in both thalamic nuclei and the STN. The differences in proportions of directional and multi-modal signals observed here in comparison with earlier reports at higher cortical structures, underscore the central role of the thalamus and STN in the regulatory feedback loops involved in the control of heart function.

### Methodological considerations

A challenge in evaluating neural activity relative to the cardiac cycle is the presence of heart cycle-related artifacts in the recorded signals (Kern et al., 2013). Extensive work on cardiac movement of the brain, using imaging approaches (e.g. Holdsworth et al., 2016; Terem et al., 2021), has highlighted the importance of carefully considering this issue when studying neural responses to the heart. MER is known to be affected by the cardioballistic effect caused by pulsating vessels near the recording electrode and observed in the waxing and waning of the extracellular signal (MacIver et al., 2011; Montgomery, Jr, 2014). This effect can affect spike detection and sorting and generate spurious modulation of the neural activity (Mosher et al., 2020). Recent work has proposed an interesting approach of post-hoc motion correction for both LFP and single unit using high density recording probes in cortical regions (Paulk et al., 2022). To overcome this major issue, given the limitations of intraoperative MER (acute single channel recordings), we opted for an approach of detecting and excluding neurons showing signs of motion. We applied methodology derived from recent work on heartbeat-related modulation of EAPs (Mosher et al., 2020) to identify neurons showing cyclical modulation of EAPs with the cardiac cycle (a sign that the neuron was possibly affected by the cardioballistic artifact) and excluded them from our sample. Accordingly, we included only neurons without significant EAP shape modulation along the cardiac cycle in further analyses (73 out of the 127 recorded units). This conservative approach aimed at ensuring that spurious modulations of the neural activity were not falsely counted as interoceptive responses. Importantly, we observe that the neurons detected here encoded different modalities (heartbeat, respiration and/or IBI), showing that the observed effects cannot relate to the cardioballistic artifact. This is further confirmed in the present data by showing various types of responses, such as the increase or decrease in firing rates, the positive and negative correlations between firing rate and IBI, as well as the positive or negative PR, and the encoding of different phases of the respiratory cycle. Collectively, these findings exclude that the present changes in neural firing in STN, Vc, and Vim were merely caused by the cardioballistic artifact.

### Respiratory signals

Analyzing the thalamic and sub-thalamic responses with respect to the cyclic respiratory signal, we were able to identify 22 neurons (∼30%) the firing activity of which was modulated by the phase of the respiratory cycle. We also report that the firings were centered around different phases of the respiratory cycle for the different neurons, with some neurons firing more likely during inspiration and others preferring expiration. To the best of our knowledge, the existence of respiratory signals in thalamic and subthalamic neurons has not been reported in humans before, but confirms previous animal data, reported from the thalamus and STN of the cat (Chen et al., 1992; Vibert et al., 1979), with about 15% of units found to correlate with the level of the respiratory drive. Corollary discharges from the brainstem respiratory neurons to the sensory cortex, could be at the basis of the thalamic signals we observed. In particular, the finding that the preferred respiratory phases vary across neurons is reminiscent of respiratory neurons in the brainstem whose activation is linked to different phases of either inspiration or expiration, controlling the generation of respiratory rhythms (Feher, 2017; Onimaru et al., 1997). Alternatively, some of the observed modulations could also be linked to mechanoreceptors in the chest wall whose projections appear to play a role in shaping respiratory sensations (Homma et al., 1988). Finally, basal ganglia and thalamus activity have also been common findings in neuroimaging studies of volitional breathing as well as investigating complex behavioral breathing acts, such as breathing during speech, while singing or exercising (Betka et al., 2022; McKay et al., 2003; Murphy et al., 2017; K. Pattinson et al., 2009; K. T. S. Pattinson et al., 2009).

We observed neurons modulated by respiratory phase in all three recorded regions, without clear differences in either proportion of responsive units or in phase preference across the three recording sites. Accordingly, recent evidence from mice has shown widespread neural responses to the breathing phase in several brain regions, including the sensory and midline thalamus (Karalis & Sirota, 2022). The breathing signal has been hypothesized to act as an oscillatory pacemaker improving functional coordination between brain regions by synchronizing neural activity (Ito et al., 2014; Karalis & Sirota, 2022). The present data show that thalamic and STN neurons may be part of this larger network of neurons involved in the coordination of brain activity. Concerning overall methodology, we also note that, given the much slower frequency of the respiratory cycle, these signals should not be affected by eventual cardiac-related artifacts or signals. Furthermore, since different neurons showed different phase preferences the present data do not appear to be linked to physiological changes during a specific phase of respiration. Additionally, a good portion of the respiration-entrained units (∼40%) was found to only encode respiratory signals (no response to the heartbeat or to the IBI), thus providing further evidence that this effect is not the result of indirect cardiac modulation.

To summarize, our results show that information about the respiratory cycle is encoded in about one third of the thalamic and subthalamic neurons. The preferred spiking phases varied across neurons, suggesting that breathing-related signals in thalamic nuclei and STN may function as oscillatory pacemaker improving functional coordination between brain regions by synchronizing neural activity.

### Thalamic and subthalamic relays of visceral signals

In this study we identified neurons in the thalamus and STN encoding different visceral information that is related to cardiac and respiratory signals. The modulation of human thalamic neuronal activity with visceral signals is compatible with several anatomical and functional studies in animals that have demonstrated widespread connectivity between the thalamus and important visceral and interoceptive centers such as the insular cortex (Cechetto & Saper, 1987; Guldin & Markowitsch, 1984; Mufson & Mesulam, 1984). Similarly, projections from STN to key structures involved in visceral processing such as the anterior cingulate cortex, the somatosensory cortex, and the insular cortex have been extensively described in rodents and primates (Afsharpour, 1985; Canteras et al., 1990; Carpenter et al., 1981; Jürgens, 1984; Kitai & Deniau, 1981; Künzle, 1977, 1978; Monakow et al., 1978; Rinvik & Ottersen, 1993; Takada et al., 2001).

Most of the neurons we identified (66%) responded to a single type of signal (either heartbeat, IBI, or respiration) and these responsive units were localized in all three investigated subcortical regions of thalamus and STN, with no clear bias in their distribution across regions. The remaining 34% of the units were found to encode two (30%), or only very rarely all three measured modalities (4%). To our knowledge, this is the first evidence of multimodal visceral neurons in the human thalamus and STN. This is consistent with data from neurons in the thalamus of non-human primates, which have shown to widely encode visceral signals (more than 80% of units responded to gastro-intestinal or bladder-related signals), with about 70% of those units encoding multimodal signals (somatic and multivisceral) (Bruggemann et al., 1994). The present evidence for multimodal visceral information at the single neuron level and at the level of each tested subcortical region suggests that, in addition to their function as a relay to efferent and afferent information to cortical regions, the thalamus and the STN also integrate different visceral inputs and outputs. Importantly, we note that the functions of heart and lung that we analyzed show physiologically close interactions, as illustrated for instance during inspiration-induced cardio-acceleration (referred to as respiratory sinus arrhythmia (Pinsky, 2006)). Together with previous evidence of widespread visceral signals encoded in subcortical regions, our findings underscore the interplay between different visceral signals and between cardiac and respiratory function in particular. These present data also reveal that visceral cardiorespiratory signals are processed in all three subcortical regions, at the level of single neurons. Many of the observed neurons, encoded more than one signal at the same time, highlighting the continuous interaction between diverse visceral information at the subcortical level, essential for successful homeostasis and behavioral control.

Overall, the present results demonstrate the direct functional involvement of thalamic and subthalamic single neurons in the processing of visceral signals in humans. The large number of detected units, the richness and diversity of signals encoded corroborate the idea that these regions are not only essential relays of the diverse visceral information reaching the brain, but are furthermore involved in regulatory feedback loops that control visceral functions. Our findings can help to better characterize the functional pathway of visceral signal processing through the subcortical regions which is of key importance also in several medical specializations (cardiology, pulmonology, neurology, psychiatry), but also psychological research. For example, the importance of the relationship between the nervous autonomic system and cardiovascular conditions, such as lethal arrhythmia, has become increasingly clear in the past decades (Task Force, 1996), and there is increasing evidence of the role of the thalamus in autonomic regulation and disturbance (Benarroch & Stotz-Potter, 2006). Visceral processing is also recognized to affect cognition and to be an important feature of some psychiatric disorders (Critchley & Garfinkel, 2017; Murphy et al., 2017). Direct and indirect measures of visceral signals and their neural correlates could in the future become important biomarkers of the state of the autonomic system, and our methodologies could support the exploration of those signals at the level of single cells. The development of chronically recording DBS implants and of novel closed loop brain stimulation interfaces also represents an ideal scenario where our investigations could find both scientific and therapeutic application.

### Limitations

While we tried to address several methodological issues, our study is still limited by the nature of MER. First of all, we are aware that all our subjects are patients suffering from either PD or essential tremor, conditions that are known to affect neural computation in subcortical regions and possibly physiological parameters (Heimrich et al., 2021). Therefore, it is possible that these diseases affected the present responses. Furthermore, it is important to note that different subcortical regions were targeted depending on disease (PD-STN; essential tremor-thalamus). The recording of neural responses in other subcortical regions involved in interoceptive processing (e.g. nucleus of the solitary tract) was not possible in the present study. Furthermore, recordings from regions presumably not involved in interoceptive processing would have been an interesting additional control. While different regions were recorded in the different patient populations, we note, however, that we observed only minor differences in the distribution of responsive units and direction of IBI correlation, while we found no difference in the latency of responses to the heartbeat and preference of respiratory phase across the different regions and hence observed similar findings whether recordings were carried out in patients with essential tremor of PD. The lack of statistically significant differences across regions could be partially explained by the variability of neural responses to interoceptive signals within each region and by the relatively small number of responsive units within each region (we may be underpowered in statistically assessing the characteristics and proportions of responses across regions). In addition, while patients are awake during MER, residual effects of anesthesia on cardio-respiratory activity cannot be completely excluded. Monitoring of several physiological parameters, such as direct recording of air flow, and increased variability of cardio-respiratory functions will be important for future investigation but seems difficult to achieve under the current OR practice for DBS implantation. Finally, further investigations should try to disentangle to possible underlying mechanisms of the various neuronal responses to the visceral activity we observed.

## Acknowledgments

The authors thank all patients for their participation and Ohio State University Medical Center staff for technical assistance. This work was supported by two donors advised by CARIGEST SA (Fondazione Teofilo Rossi di Montelera e di Premuda and a second one wishing to remain anonymous) to O.B. We thank Prof. Catherine Tallon-Baudry (Ecole Normale Supérieure, Paris, France) for the insightful discussion and early feedback on the manuscript.

## Author contributions

Conceptualization: O.B. and M.S.; Methodology: E.D.F. and M.S.; Formal Analysis, Software, Data curation: E.D.F.; contribution to data analysis: M.S, F.B., M.B; Data collection: M.S. and N.Y.; V.K. and A.R. performed the surgeries; writing – original draft: E.D.F., M.S. and O.B.; writing – review and editing: all authors.

## Methods

### Patients and recordings

We acquired intraoperative microelectrode recordings (MER) from 23 patients (9 females, mean age: 64 years (SD: 6.0 years), Body Mass Index range 20-35kg/m2) undergoing surgical implantation of DBS electrodes in the subthalamic nucleus and thalamus for treatment of refractory Parkinson disease (N=9) or essential tremor (N=14), respectively. Patients with untreated depression or severe anxiety were not recruited. Recordings were performed at Ohio State University Medical Center between March 2017 and July 2018. All patients signed informed consent to participate in the study before surgery. Experimental protocol was approved by the Ohio State University Medical Center Institutional Review Board (Columbus, Ohio).

MERs during DBS surgery are used to accurately localize the target region (i.e. the dorsolateral part of the subthalamic nucleus, STN, for Parkinson treatment and the Ventral Intermedius thalamic nuclei (Vim) for essential tremor). MERs from Ventral caudalis (Vc) thalamic nuclei are typically used to determine the boundaries of the sensory-motor region of the thalamus thus improving Vim localization (Marks, 2015). During each intraoperative session, several recording blocks were acquired from the STN, Vim, and/or Vc. All recordings were collected at the beginning of the surgery before any stimulation or motor task. Patients were awake with blood pressure kept below 140 mm Hg and did not receive specific instruction, except to avoid any voluntary movement and to keep their eyes open. Dexmedetomidine was used at the beginning of the surgery and interrupted during the burr hole to ensure a 30’ washout period prior to MER recordings. We present data from 63 recording chunks with at least one neuron (22 from STN, 18 from Vc, 23 from Vim), the average duration of those recording blocks was 148s (SD: 43.3s, Range 52.8s to 273.4s), with an average of 202 cardiac cycles recorded (SD=71, range 60-373). Electrode locations were based solely on medical need and were calculated using stereotactic coordinates extracted from preoperative MRI. However, we note that 3D inaccuracy, pneumocephalus or image fusion errors induce implantation imprecision, and some caution is necessary regarding the exact recording location during DBS. Electrophysiological recordings were acquired with a sampling rate of 48 kHz using the FHC 4000 LP (Lead Positioner) Neuromodulation Targeting System (manufactured by FHC, Inc., 1201 Main St., Bowdoin, ME USA 04287). Simultaneous acquisition of electrocardiogram (ECG) was performed by means of two surface electrodes placed on the patient chest and a reference electrode placed on the right ankle. ECG was initially sampled at 48 kHz.

### Signal processing and spike extraction

All recorded signals were processed offline using Matlab. R-peaks were detected from the ECG signal with the HEPLAB package (Perakakis, 2019) using the ‘ecglab fast’ algorithm for peak detections (de Carvalho et al., 2002). Detected R-peaks were then inspected manually to exclude spurious events. Aberrant cardiac cycles with IBI <0.5 or IBI>1.5 (corresponding, respectively to a cardiac rate >120bmp and <40bpm) were excluded from all further computations. The heart rate variability was assessed from the standard deviation of the IBIs for all beats (SDRR, equivalent to SDNN). The distribution of IBIs, heart rates, and SDRRs across recording chunks is shown in Figure S2. Electrophysiological signals were processed using the latest implementation of Wave-clus (Chaure et al., 2018) to extract and sort single neuron activity. Signal was initially high-pass filtered (above 300Hz), single spike activity was then extracted using a detection threshold equal to 5 times the estimated standard deviation of the background noise (*σ_N_*). Spikes from different putative neurons were sorted using the automatic implementation of Wave-clus. Initial spike sorting was then reviewed manually to exclude noise clusters and optimize cluster separation. Signal-to-noise ratio (SNR) for each identified neuron was computed as the mean amplitude of each neuron’s waveforms over the background noise. SNR distribution is shown in Figure S3-A (mean SNR = 9.86 (SD=6.3). Neurons with more than 3% of refectory violations (Inter-Spike interval, ISI, < 3ms) or not enough spikes (mean firing rate <0.1Hz) were excluded. The distribution of percentage of refractory violations is shown in Figure S3-B. Isolation distance (Harris et al., 2001)(Harris et al., 2001), a measure of the quality of a cluster separation, was also computed for each identified single neuron based on the top 10 high-dimensional spike features extracted during sorting. Its distribution is reported in Figure S3-C (note that the Isolation Distance is only defined for cases in which the number of spikes outside the cluster is greater than the number of spikes in the cluster)

### EAP waveform modulation test

One of the effects of cardioballistic artifacts is the cyclical change in the shape of Extracellular Action potential (EAP) from the recorded neurons. Stability of the Extracellular Action potential waveform was tested using an implementation based on a recently proposed approach (Mosher et al., 2020). The methodology aims at assessing the degree of variation of several EAP features along the cardiac cycle. The EAP features we measured were: maximum amplitude (AMP) in μV measured at the trough time; half-width time (HW) in ms corresponding to the total time that the EAP trough is below half AMP; trough-to-peak time (TPW) in ms corresponding to the time between the trough and the following peak; time for repolarization (REP) in ms measuring the time after peak when the EAP reaches half peak value (see Figure S1A). The test was performed as follows. From the high-pass filtered signal at 32kHz, the EAPs were sampled around the detection peak taking 25 samples (0.78ms) before and 57 (1.78ms) samples after peak. Samples were then interpolated to up-sampled the EAP to 256 samples (100 kHz) and realigned to the EAP trough (see Figure S1 panels A1 and B1). Changes in the EAP features throughout the cardiac cycle were then assessed by binning the spikes in 100ms bins (with 10ms overlap) around the R-peak and calculating an average EAP waveform for each bin. Bins started 100ms before R-peak and ended at the ‘low IBI’ value (corresponding to the 2.5th percentile of all IBI durations for the given recording). From the average waveforms, EAP features were extracted and evaluated along the cardiac cycle. The degree of change along the cycle was evaluated by fitting a cosine function to the data points using circular linear-regression. For this purpose, the selected time window was transformed into radians (with 0ms corresponding to 0rad, and the maximum time corresponding to 2π rad) to represent the angular phase of the cardiac cycle phase. The amplitude of the fitted function indicates the maximum degree of modulation of the feature in percentage and is termed Modulation Index (MI) (see Figure S1 panels A3 and B3). A feature was considered to be significantly modulated by the cardiac cycle if: the model indicated that the amplitude of the fit was significantly different from zero (t-test, p<0.05) and significantly larger than a bootstrapped null statistic (p<0.05). The null statistic was obtained by bootstrapping the data 1000 times using surrogate R-peaks which preserved the same IBI and variability as the original R-peaks but were randomly shifted in time with a maximum absolute shift of 500ms (Park et al., 2014, 2018). EAP waveforms that had at least one EAP feature significantly modulated by the cardiac cycle were labeled as modulated and those neurons were excluded from further analysis.

### Neural responses to heartbeat

Modulation of the neural firing rates in response to the heartbeat was tested for each neuron as follows. For each detected R-peak, we considered all neural spikes falling in a time interval around the time of the R-peak. The time interval ranged from -100ms to the median value of the cardiac cycle duration (median IBI) for the given recording. Spike times were realigned relatively to the time of the R-peak and binned into 10 equally spaced bins. The time bins were then converted into radians to represent the angular phase along the cardiac cycle (with 0ms corresponding to 0rad, and the maximum time corresponding to 2π rad). We then computed the z-statistic for the non-uniformity of circular data (Rayleigh’s test) on the binned spikes and compared it to a null distribution of 1000 surrogate statistics obtained by jittering the R-peaks (between -500ms and +500ms) while preserving the IBI and variability of cardiac cycles (Park et al., 2014, 2018). In order to ensure a robust estimation of modulation across trials, we additionally measured the spike-field coherence between the neural spikes and the phase of the cardiac cycle (averaged in the frequency band < 20Hz). For a neuron to have a significant modulation of firing rates in response to the heartbeat we asked that the z-statistic for non-uniformity was larger than 95% of the surrogates (p<0.05) and that the coherence between spikes and cardiac phase was larger than the coherence significance level (Thompson, 1979):

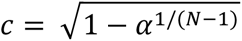

with α alpha equal to 0.05 significance level corrected for multiple comparisons (Bonferroni correction) and N being the number of cardiac cycles used to measure coherence. Additionally, we repeated the response analysis but this time time-locking the spikes to the end of the cardiac cycle. If the neural response was disrupted when time-locked to the end of the cycle, this would disambiguate the directionality of the effect (from heart to neuron or vice versa).

For neurons with a significant modulation of firing rate along the cardiac cycle we also calculated the peak latency of the firing rate change. To do so, we first calculated the average instantaneous firing rate along the cardiac cycle by convolving the spikes with a Gaussian kernel with σ=10ms (truncated at 1% amplitude) and then averaging across all cardiac cycles. Given that some neurons responded with an increase in activity, while others responded with a suppression in activity along the cardiac cycle, we first identified the direction of change in the time interval following the R-peak (from 50ms to 400ms after R-peak, the sign of the slope of a linear fit provided the direction of change). The peak latency in ms was then extracted as the time where the firing rate was maximum/minimum for increased/suppressed activation respectively. Finally, the peak latency was expressed in radians along the cardiac cycle (with R-peaks corresponding to 0 and 2π rad) by normalizing the latency value in s by the average IBI measured during the session. This was done in order to obtain latency values comparable across the different recording sessions.

### Relationship between spike firing and Inter-beat Intervals

We assessed the relationship between neural firing and duration of the cardiac cycle as follows. For each cardiac cycle we measured the Interbeat Interval (IBI) which corresponds to the time between one ECG R-peak and the following one. For each neuron, the average firing rate during each cardiac cycle (FRc, measured in Hz) was calculated as the number of spikes fired during the cycle divided by the corresponding IBI. Correlation between FRc and IBI was calculated using Pearson’s correlation. Statistical significance of the correlation was assessed from t-distributions and corrected for multiple comparisons (Bonferroni correction). In order to test for a directional effect from the spiking activity to the IBI we used a phase-response (*PR*) analysis, as recently proposed by Kim et al. (K. Kim et al., 2019). PR analysis measures how much the cardiac cycle is lengthened or shortened in response to spiking activity as follows. For each spike *i*, the phase-response *PR_i_* is obtained by comparing the time from the spike to the following R-peak (*Tpost_i_*) with its expected value of *TpostExp_i_*:

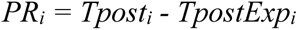

With the expected value calculated from:

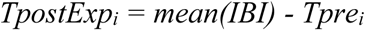

Where *mean(IBI)* is the mean IBI for the recording session and *Tpre_i_* is the time from the considered spike to the previous R-peak. For each neuron, a unique value for the PR is obtained as the mean of *PR_i_* on all spikes fired. Significance of PR was assessed by comparing the PR value with a surrogate distribution of 10000 PR values obtained by jittering the spike times (between -500ms and +500ms). The p-values obtained were corrected for multiple comparisons using Bonferroni correction.

### ECG-derived respiration extraction

Since we had no direct recording of respiratory signals due to surgical constraints, we extracted a putative respiratory signal from the ECG traces using an ECG-Derived Respiration (EDR) algorithm. The ECG signal is secondarily modulated by respiration because of the cyclic change in the positions of ECG electrodes on the chest surface relative to the heart as the lungs fill and empty. This movement changes the impedance between the heart and the ECG leads and generates a modulation in the recorded ECG signal. The EDR is a well-known extraction technique (Moody et al., 1985) that has been further developed and evaluated in recent years (Charlton et al., 2016). In our dataset, we extracted the EDR using a feature-based extraction algorithm implemented in the RRest toolbox (http://peterhcharlton.github.io/RRest) (Charlton et al., 2016). Feature-based extraction methods consist in measuring a beat-by-beat feature whose changes in time reflect the respiratory signal. We considered 3 different features in our analysis: amplitude modulation (AM, estimated as difference between amplitudes of troughs and proceeding R-peaks), frequency modulation (FM, estimated as the time interval between consecutive R-peaks), and baseline wander (BW, estimated as mean amplitude of troughs and proceeding R-peaks). The algorithm includes several steps briefly detailed here (please refer to the original publication for further details): From the raw ECG signal, high frequencies (over 100 Hz) are eliminated by low-pass filtering; R-peaks are detected and fiducial points of the ECG QRS complex are identified; the target features of the signal are then measured (AM, FM, and BW. The extracted features are then resampled at regular sampling frequency (5 Hz) and finally very low frequencies are eliminated (below 0.07Hz corresponding to 4 breaths per minute (bpm)). In order to assess the quality of the ECG-derived respiratory signal, we performed the quality control evaluation proposed by Karlen et al. (Karlen et al., 2013). For each recording segment (10s), the respiratory rate was estimated using the Count-Origin algorithm (Schäfer & Kratky, 2008) for each of the three features. If there was no agreement between the three methods (SD between the three extracted rates > 4bpm), the segment was considered aberrant and excluded from further analyses. For all the accepted segments, the respiratory signals extracted from AM were saved and used in the following analyses. For almost all sessions (62 out 63) we were able to correctly extract at least one good EDR segment. The mean respiratory rate measure across sessions was 17.7 breaths per minute (SD = 1.7), a full distribution of respiratory rates across recording blocks is shown in Figure S7-B.

### EAP waveform along respiratory cycle

While the cardioballistic artifact is the most prominent artifact observed in intracranial recordings, it is possible that additional artifacts caused by pressure changes linked to the respiratory cycle could be present as well. To exclude any additional contamination in our data we adapted the EAP waveform modulation test to verify whether any cyclical change in EAP was observable during the respiratory cycle. The test was implemented similarly to the cardiac one, by extracting the EAPs features along respiratory cycles aligned to the beginning of inspiration and comparing the measured motion index to those extracted from a surrogate population. Among the 127 neurons initially identified, we found that 7 units exhibited a significant EAP modulation linked to the respiratory cycle. However, all 7 of them belonged to the population of neurons already excluded based on the cardiac EAP modulation. Thus, none of the 73 neurons included in our analysis exhibited a significant EAP modulation linked to the respiratory cycle.

### Neural responses to respiration

The phase of the respiratory signal was extracted as the angular component of the Hilbert transform of the HDR signals. For each neural spike, the corresponding respiratory phase (spike-phase) was extracted as the instantaneous respiratory phase at the time of the spike. Non-uniformity of the spike density phase distribution was tested using a Rayleigh’s test for circular data. Significance of the modulation was assessed by comparing the z-statistic obtained with a surrogate statistic (10000 surrogates) obtained by jittering the spike times (between -2.5s and +2.5s). Note that in this case a large interval for jittering was chosen in order to break time dependency between the spikes and the slow respiratory rhythm.

### Codes and data availability

Codes for analysis were custom made in Matlab and the following toolboxes were used: EAP waveforms were tested using partially the code from Mosher et al. (Mosher et al., 2020); R-peaks were detected with the HEPLAB package (Perakakis, 2019); RRest toolbox for ECG-derived respiration (http://peterhcharlton.github.io/RRest) (Charlton et al., 2016).

Analysis scripts will be available in a public repository following publication. After publication, neural data will be available upon written request, dependent upon the establishment of a data sharing agreement between the trial’s investigator, sponsor, and interested third party.

## References

Adler, D., Herbelin, B., Similowski, T., & Blanke, O. (2014). Breathing and sense of self: Visuo–respiratory conflicts alter body self-consciousness. Respiratory Physiology & Neurobiology, 203, 68–74. 10.1016/j.resp.2014.08.003

Afsharpour, S. (1985). Topographical projections of the cerebral cortex to the subthalamic nucleus. The Journal of Comparative Neurology, 236(1), 14–28. 10.1002/cne.902360103

Allard, E., Canzoneri, E., Adler, D., Morélot-Panzini, C., Bello-Ruiz, J., Herbelin, B., Blanke, O., & Similowski, T. (2017). Interferences between breathing, experimental dyspnoea and bodily self-consciousness. Scientific Reports, 7(1), 9990. 10.1038/s41598-017-11045-y

Armour, J., & Ardell, J. (2004). Basic and clinical neurocardiology. Oxford University Press.

Asato, F., & Yokota, T. (1989). Responses of neurons in nucleus ventralis posterolateralis of the cat thalamus to hypogastric inputs. Brain Research, 488(1–2), 135–142. 10.1016/0006-8993(89)90702-6

Beissner, F., Meissner, K., Bar, K.-J., & Napadow, V. (2013). The Autonomic Brain: An Activation Likelihood Estimation Meta-Analysis for Central Processing of Autonomic Function. Journal of Neuroscience, 33(25), 10503–10511. 10.1523/JNEUROSCI.1103-13.2013

Benarroch, E. E., & Stotz-Potter, E. H. (2006). Dysautonomia in Fatal Familial Insomnia as an Indicator of the Potential Role of the Thalamus in Autonomie Control. Brain Pathology, 8(3), 527–530. 10.1111/j.1750-3639.1998.tb00174.x

Berntson, G. G., Cacioppo, J. T., & Quigley, K. S. (1993). Cardiac psychophysiology and autonomic space in humans: Empirical perspectives and conceptual implications. Psychological Bulletin, 114(2), 296–322. 10.1037/0033-2909.114.2.296

Berntson, G. G., & Khalsa, S. S. (2021). Neural Circuits of Interoception. Trends in Neurosciences, 44(1), 17–28. 10.1016/j.tins.2020.09.011

Berntson, G. G., Quigley, K. S., & Lozano, D. (2007). Cardiovascular Psychophysiology. In G. Berntson, J. T. Cacioppo, & L. G. Tassinary (Eds.), Handbook of Psychophysiology (3rd ed., pp. 182–210). Cambridge University Press. https://www.cambridge.org/core/books/handbook-of-psychophysiology/cardiovascular-psychophysiology/CAB9D08704751D5A26A58C442B3F2BB8

Betka, S., Adler, D., Similowski, T., & Blanke, O. (2022). Breathing control, brain, and bodily self-consciousness: Toward immersive digiceuticals to alleviate respiratory suffering. Biological Psychology, 171, 108329. 10.1016/j.biopsycho.2022.108329

Birznieks, I., Boonstra, T. W., & Macefield, V. G. (2012). Modulation of Human Muscle Spindle Discharge by Arterial Pulsations—Functional Effects and Consequences. PLoS ONE, 7(4), e35091. 10.1371/journal.pone.0035091

Bishop, V. S., Malliani, A., & Thorén, P. (2011). Cardiac Mechanoreceptors. In Comprehensive Physiology (pp. 497–555). John Wiley & Sons, Ltd. 10.1002/cphy.cp020315

Blanke, O., Slater, M., & Serino, A. (2015). Behavioral, Neural, and Computational Principles of Bodily Self-Consciousness. Neuron, 88(1), 145–166. 10.1016/j.neuron.2015.09.029

Bruggemann, J., Shi, T., & Apkarian, A. (1994). Squirrel monkey lateral thalamus. II. Viscerosomatic convergent representation of urinary bladder, colon, and esophagus. The Journal of Neuroscience, 14(11), 6796–6814. 10.1523/JNEUROSCI.14-11-06796.1994

Canteras, N. S., Shammah-Lagnado, S. J., Silva, B. A., & Ricardo, J. A. (1990). Afferent connections of the subthalamic nucleus: A combined retrograde and anterograde horseradish peroxidase study in the rat. Brain Research, 513(1), 43–59. 10.1016/0006-8993(90)91087-W

Carpenter, M. B., Carleton, S. C., Keller, J. T., & Conte, P. (1981). Connections of the subthalamic nucleus in the monkey. Brain Research, 224(1), 1–29. 10.1016/0006-8993(81)91113-6

Cechetto, D. F. (2014). Cortical control of the autonomic nervous system: Cortical autonomic control. Experimental Physiology, 99(2), 326–331. 10.1113/expphysiol.2013.075192

Cechetto, D. F., & Saper, C. B. (1987). Evidence for a viscerotopic sensory representation in the cortex and thalamus in the rat. The Journal of Comparative Neurology, 262(1), 27–45. 10.1002/cne.902620104

Chandler, M. J., Hobbs, S. F., Fu, Q.-G., Kenshalo, D. R., Blair, R. W., & Foreman, R. D. (1992). Responses of neurons in ventroposterolateral nucleus of primate thalamus to urinary bladder distension. Brain Research, 571(1), 26–34. 10.1016/0006-8993(92)90506-5

Charlton, P. H., Bonnici, T., Tarassenko, L., Clifton, D. A., Beale, R., & Watkinson, P. J. (2016). An assessment of algorithms to estimate respiratory rate from the electrocardiogram and photoplethysmogram. Physiological Measurement, 37(4), 610–626. 10.1088/0967-3334/37/4/610

Chaure, F. J., Rey, H. G., & Quian Quiroga, R. (2018). A novel and fully automatic spike-sorting implementation with variable number of features. Journal of Neurophysiology, 120(4), 1859–1871. 10.1152/jn.00339.2018

Chen, Z., Eldridge, F. L., & Wagner, P. G. (1992). Respiratory-associated thalamic activity is related to level of respiratory drive. Respiration Physiology, 90(1), 99–113. 10.1016/0034-5687(92)90137-L

Coenen, V. A., Gielen, F. L. H., Castro-Prado, F., Abdel Rahman, A., & Honey, C. R. (2008). Noradrenergic modulation of subthalamic nucleus activity in human: Metoprolol reduces spiking activity in microelectrode recordings during deep brain stimulation surgery for Parkinson’s disease. Acta Neurochirurgica, 150(8), 757–762. 10.1007/s00701-008-1619-5

Craig, A. (2009). How do you feel — now? The anterior insula and human awareness. Nature Reviews Neuroscience, 10(1), 59–70. 10.1038/nrn2555

Critchley, H. D., & Garfinkel, S. N. (2017). Interoception and emotion. Current Opinion in Psychology, 17, 7–14. 10.1016/j.copsyc.2017.04.020\

Critchley, H. D., & Harrison, N. A. (2013). Visceral Influences on Brain and Behavior. Neuron, 77(4), 624–638. 10.1016/j.neuron.2013.02.008

Damasio, A., & Carvalho, G. B. (2013). The nature of feelings: Evolutionary and neurobiological origins. Nature Reviews Neuroscience, 14(2), 143–152. 10.1038/nrn3403

de Carvalho, J. L. A., da Rocha, A. F., de Oliveira Nascimento, F. A., Neto, J. S., & Junqueira, L. F. (2002). Development of a Matlab software for analysis of heart rate variability. 6th International Conference on Signal Processing, 2002., 2, 1488–1491. 10.1109/ICOSP.2002.1180076

Dlouhy, B. J., Gehlbach, B. K., Kreple, C. J., Kawasaki, H., Oya, H., Buzza, C., Granner, M. A., Welsh, M. J., Howard, M. A., Wemmie, J. A., & Richerson, G. B. (2015). Breathing Inhibited When Seizures Spread to the Amygdala and upon Amygdala Stimulation. Journal of Neuroscience, 35(28), 10281–10289. 10.1523/JNEUROSCI.0888-15.2015

Dulla, C. G., Dobelis, P., Pearson, T., Frenguelli, B. G., Staley, K. J., & Masino, S. A. (2005). Adenosine and ATP Link PCO2 to Cortical Excitability via pH. Neuron, 48(6), 1011– 1023. 10.1016/j.neuron.2005.11.009

Eldridge, F. L., Millhorn, D. E., Killey, J. P., & Waldrop, T. G. (1985). Stimulation by central command of locomotion, respiration and circulation during exercise. Respiration Physiology, 59(3), 313–337. 10.1016/0034-5687(85)90136-7

Eldridge, F. L., Millhorn, D. E., & Waldrop, T. G. (1981). Exercise Hyperpnea and Locomotion: Parallel Activation from the Hypothalamus. Science, 211(4484), 844– 846. 10.1126/science.7466362

Emmers, R. (1966). Modulation of the thalamic relay of taste by stimulation of the tongue with ice water. Experimental Neurology, 16(1), 50–56. 10.1016/0014-4886(66)90085-9

Engelen, T., Solcà, M., & Tallon-Baudry, C. (2023). Interoceptive rhythms in the brain. Nature Neuroscience. 10.1038/s41593-023-01425-1

Faull, O. K., Jenkinson, M., Clare, S., & Pattinson, K. T. S. (2015). Functional subdivision of the human periaqueductal grey in respiratory control using 7tesla fMRI. NeuroImage, 113, 356–364. 10.1016/j.neuroimage.2015.02.026

Feher, J. (2017). Control of Ventilation. In Quantitative Human Physiology (pp. 672–681). Elsevier. 10.1016/B978-0-12-800883-6.00066-5

Ford, T. W., & Kirkwood, P. A. (2018). Cardiac modulation of alpha motoneuron discharges. Journal of Neurophysiology, 119(5), 1723–1730. 10.1152/jn.00025.2018

Frommer, G. P. (1961). Gustatory afferent responses in the thalamus. In M. R. Kare & B. P. Halpern The physiological and behavioral aspects of taste (pp. 50–59). Univer. Chicago Press.

Frysinger, R. C., & Harper, R. M. (1990). Cardiac and Respiratory Correlations with Unit Discharge in Epileptic Human Temporal Lobe. Epilepsia, 31(2), 162–171. 10.1111/j.1528-1167.1990.tb06301.x

Gray, M. A., Taggart, P., Sutton, P. M., Groves, D., Holdright, D. R., Bradbury, D., Brull, D., & Critchley, H. D. (2007). A cortical potential reflecting cardiac function. Proceedings of the National Academy of Sciences, 104(16), 6818–6823. 10.1073/pnas.0609509104

Green, A. L., & Paterson, D. J. (2020). Using Deep Brain Stimulation to Unravel the Mysteries of Cardiorespiratory Control. In R. Terjung (Ed.), Comprehensive Physiology (1st ed., pp. 1085–1104). Wiley. 10.1002/cphy.c190039

Groenewegen, H. J., & Witter, M. P. (2004). Thalamus. In The Rat Nervous System (pp. 407–453). Elsevier. 10.1016/B978-012547638-6/50018-3

Guldin, W. O., & Markowitsch, H. J. (1984). Cortical and thalamic afferent connections of the insular and adjacent cortex of the cat. The Journal of Comparative Neurology, 229(3), 393–418. 10.1002/cne.902290309

Haines, D. E., & Mihailoff, G. A. (2018). Fundamental neuroscience for basic and clinical applications (5th ed.). http://liverpool.idm.oclc.org/login?url=https://www.clinicalkey.com/dura/browse/bookChapter/3-s2.0-C20140037185

Harris, K. D., Hirase, H., Leinekugel, X., Henze, D. A., & Buzsáki, G. (2001). Temporal Interaction between Single Spikes and Complex Spike Bursts in Hippocampal Pyramidal Cells. Neuron, 32(1), 141–149. 10.1016/S0896-6273(01)00447-0

Hayward, L. F., & Felder, R. B. (1995). Cardiac rhythmicity among NTS neurons and its relationship to sympathetic outflow in rabbits. American Journal of Physiology-Heart and Circulatory Physiology, 269(3), H923–H933. 10.1152/ajpheart.1995.269.3.H923

Heimrich, K. G., Lehmann, T., Schlattmann, P., & Prell, T. (2021). Heart Rate Variability Analyses in Parkinson’s Disease: A Systematic Review and Meta-Analysis. Brain Sciences, 11(8), 959. 10.3390/brainsci11080959

Herrero, J. L., Khuvis, S., Yeagle, E., Cerf, M., & Mehta, A. D. (2018). Breathing above the brain stem: Volitional control and attentional modulation in humans. Journal of Neurophysiology, 119(1), 145–159. 10.1152/jn.00551.2017

Hirai, T., & Jones, E. G. (1989). A new parcellation of the human thalamus on the basis of histochemical staining. Brain Research Reviews, 14(1), 1–34. 10.1016/0165-0173(89)90007-6

Holdsworth, S. J., Rahimi, M. S., Ni, W. W., Zaharchuk, G., & Moseley, M. E. (2016). Amplified magnetic resonance imaging (aMRI): Amplified MRI (aMRI). Magnetic Resonance in Medicine, 75(6), 2245–2254. 10.1002/mrm.26142

Homma, I., Kanamaru, A., & Sibuya, M. (1988). Proprioceptive Chest Wall Afferents and the Effect on Respiratory Sensation. In Respiratory Psychophysiology (In von Euler, C. and Katz-Solomon, M.).

Ito, J., Roy, S., Liu, Y., Cao, Y., Fletcher, M., Lu, L., Boughter, J. D., Grün, S., & Heck, D. H. (2014). Whisker barrel cortex delta oscillations and gamma power in the awake mouse are linked to respiration. Nature Communications, 5(1), 3572. 10.1038/ncomms4572

Jürgens, U. (1984). The efferent and afferent connections of the supplementary motor area. Brain Research, 300(1), 63–81. 10.1016/0006-8993(84)91341-6

Karalis, N., & Sirota, A. (2022). Breathing coordinates cortico-hippocampal dynamics in mice during offline states. Nature Communications, 13(1), 467. 10.1038/s41467-022-28090-5

Karlen, W., Raman, S., Ansermino, J. M., & Dumont, G. A. (2013). Multiparameter Respiratory Rate Estimation From the Photoplethysmogram. IEEE Transactions on Biomedical Engineering, 60(7), 1946–1953. 10.1109/TBME.2013.2246160

Kern, M., Aertsen, A., Schulze-Bonhage, A., & Ball, T. (2013). Heart cycle-related effects on event-related potentials, spectral power changes, and connectivity patterns in the human ECoG. NeuroImage, 81, 178–190. 10.1016/j.neuroimage.2013.05.042

Khalsa, S. S., Feinstein, J. S., Simmons, W. K., & Paulus, M. P. (2018). Taking Aim at Interoception’s Role in Mental Health. Biological Psychiatry: Cognitive Neuroscience and Neuroimaging, 3(6), 496–498. 10.1016/j.bpsc.2018.04.007

Khalsa, S. S., & Lapidus, R. C. (2016). Can Interoception Improve the Pragmatic Search for Biomarkers in Psychiatry? Frontiers in Psychiatry, 7. 10.3389/fpsyt.2016.00121

Khan, A. A., Lip, G. Y. H., & Shantsila, A. (2019). Heart rate variability in atrial fibrillation: The balance between sympathetic and parasympathetic nervous system. European Journal of Clinical Investigation, 49(11). 10.1111/eci.13174

Kim, K. J., Ramiro Diaz, J., Iddings, J. A., & Filosa, J. A. (2016). Vasculo-Neuronal Coupling: Retrograde Vascular Communication to Brain Neurons. The Journal of Neuroscience, 36(50), 12624–12639. 10.1523/JNEUROSCI.1300-16.2016

Kim, K., Ladenbauer, J., Babo-Rebelo, M., Buot, A., Lehongre, K., Adam, C., Hasboun, D., Lambrecq, V., Navarro, V., Ostojic, S., & Tallon-Baudry, C. (2019). Resting-State Neural Firing Rate Is Linked to Cardiac-Cycle Duration in the Human Cingulate and Parahippocampal Cortices. The Journal of Neuroscience, 39(19), 3676–3686. 10.1523/JNEUROSCI.2291-18.2019

Kitai, S. T., & Deniau, J. M. (1981). Cortical inputs to the subthalamus: Intracellular analysis. Brain Research, 214(2), 411–415. 10.1016/0006-8993(81)91204-X

Kluger, D. S., Balestrieri, E., Busch, N. A., & Gross, J. (2021). Respiration aligns perception with neural excitability. ELife, 10, e70907. 10.7554/eLife.70907

Kluger, D. S., & Gross, J. (2021). Respiration modulates oscillatory neural network activity at rest. PLOS Biology, 19(11), e3001457. 10.1371/journal.pbio.3001457

Künzle, H. (1977). Projections from the primary somatosensory cortex to basal ganglia and thalamus in the monkey. Experimental Brain Research, 30(4). 10.1007/BF00237639

Künzle, H. (1978). An Autoradiographic Analysis of the Efferent Connections from Premotor and Adjacent Prefrontal Regions (Areas 6 and 9) in Macaca fascicularis. Brain, Behavior and Evolution, 15(3), 185–209. 10.1159/000123779

Lenz, F. A., Seike, M., Lin, Y. C., Baker, F. H., Rowland, L. H., Gracely, R. H., & Richardson, R. T. (1993). Neurons in the area of human thalamic nucleus ventralis caudalis respond to painful heat stimuli. Brain Research, 623(2), 235–240. 10.1016/0006-8993(93)91433-S

Macefield, V. G. (2003). Cardiovascular and Respiratory Modulation of Tactile Afferents in the Human Finger Pad. Experimental Physiology, 88(5), 617–625. 10.1113/eph8802548

MacIver, M. B., Bronte-Stewart, H. M., Henderson, J. M., Jaffe, R. A., & Brock-Utne, J. G. (2011). Human Subthalamic Neuron Spiking Exhibits Subtle Responses to Sedatives. Anesthesiology, 115(2), 254–264. 10.1097/ALN.0b013e3182217126

MacKinnon, S., Gevirtz, R., McCraty, R., & Brown, M. (2013). Utilizing Heartbeat Evoked Potentials to Identify Cardiac Regulation of Vagal Afferents During Emotion and Resonant Breathing. Applied Psychophysiology and Biofeedback, 38(4), 241–255. 10.1007/s10484-013-9226-5

Marina, N., Christie, I. N., Korsak, A., Doronin, M., Brazhe, A., Hosford, P. S., Wells, J. A., Sheikhbahaei, S., Humoud, I., Paton, J. F. R., Lythgoe, M. F., Semyanov, A., Kasparov, S., & Gourine, A. V. (2020). Astrocytes monitor cerebral perfusion and control systemic circulation to maintain brain blood flow. Nature Communications, 11(1), 131. 10.1038/s41467-019-13956-y

Marks, W. J. Jr. (Ed.). (2015). Deep Brain Stimulation Management (2nd ed.). Cambridge University Press. 10.1017/CBO9781316026625

Massimini, M., Porta, A., Mariotti, M., Malliani, A., & Montano, N. (2000). Heart rate variability is encoded in the spontaneous discharge of thalamic somatosensory neurones in cat. The Journal of Physiology, 526(2), 387–396. 10.1111/j.1469-7793.2000.t01-1-00387.x

Mazziotta, J. C., Toga, A. W., Evans, A., Fox, P., & Lancaster, J. (1995). A Probabilistic Atlas of the Human Brain: Theory and Rationale for Its Development. NeuroImage, 2(2), 89–101. 10.1006/nimg.1995.1012

Mccraty, R., & Shaffer, F. (2015). Heart Rate Variability: New Perspectives on Physiological Mechanisms, Assessment of Self-regulatory Capacity, and Health Risk. Global Advances in Health and Medicine, 4(1), 46–61. 10.7453/gahmj.2014.073

McKay, L. C., Evans, K. C., Frackowiak, R. S. J., & Corfield, D. R. (2003). Neural correlates of voluntary breathing in humans. Journal of Applied Physiology, 95(3), 1170–1178. 10.1152/japplphysiol.00641.2002

Monakow, K. H., Akert, K., & K□nzle, H. (1978). Projections of the precentral motor cortex and other cortical areas of the frontal lobe to the subthalamic nucleus in the monkey. Experimental Brain Research, 33(3–4). 10.1007/BF00235561

Montgomery, Jr, E. B. (2014). Intraoperative Neurophysiological Monitoring for Deep Brain Stimulation: Principles, Practice and Cases. Oxford University Press. 10.1093/med/9780199351008.001.0001

Montoya, P., Schandry, R., & Müller, A. (1993). Heartbeat evoked potentials (HEP): Topography and influence of cardiac awareness and focus of attention. Electroencephalography and Clinical Neurophysiology/Evoked Potentials Section, 88(3), 163–172. 10.1016/0168-5597(93)90001-6

Moody, G. B., Mark, R. G., Zoccola, A., & Mantero, S. (1985). Derivation of respiratory signals from multi-lead ECGs. Computers in Cardiology, 12(1985), 113–116.

Mosher, C. P., Mamelak, A. N., Malekmohammadi, M., Pouratian, N., & Rutishauser, U. (2021). Distinct roles of dorsal and ventral subthalamic neurons in action selection and cancellation. Neuron, 109(5), 869–881.e6. 10.1016/j.neuron.2020.12.025

Mosher, C. P., Wei, Y., Kamiński, J., Nandi, A., Mamelak, A. N., Anastassiou, C. A., & Rutishauser, U. (2020). Cellular Classes in the Human Brain Revealed In Vivo by Heartbeat-Related Modulation of the Extracellular Action Potential Waveform. Cell Reports, 30(10), 3536–3551.e6. 10.1016/j.celrep.2020.02.027

Mufson, E. J., & Mesulam, M. M. (1984). Thalamic connections of the insula in the rhesus monkey and comments on the paralimbic connectivity of the medial pulvinar nucleus. The Journal of Comparative Neurology, 227(1), 109–120. 10.1002/cne.902270112

Murphy, J., Brewer, R., Catmur, C., & Bird, G. (2017). Interoception and psychopathology: A developmental neuroscience perspective. Developmental Cognitive Neuroscience, 23, 45–56. 10.1016/j.dcn.2016.12.006

Nord, C. L., & Garfinkel, S. N. (2022). Interoceptive pathways to understand and treat mental health conditions. Trends in Cognitive Sciences, S1364661322000626. 10.1016/j.tics.2022.03.004

Ohye, C., Shibazaki, T., Hirai, T., Wada, H., Hirato, M., & Kawashima, Y. (1989). Further physiological observations on the ventralis intermedius neurons in the human thalamus. Journal of Neurophysiology, 61(3), 488–500. 10.1152/jn.1989.61.3.488

Onimaru, H., Arata, A., & Homma, I. (1997). Neuronal Mechanisms of Respiratory Rhythm Generation: An Approach Using In Vitro Preparation. The Japanese Journal of Physiology, 47(5), 385–403. 10.2170/jjphysiol.47.385

Oppenheimer, S. M., Kulshreshtha, N., Lenz, F. A., Zhang, Z., Rowland, L. H., & Dougherty, P. M. (1998). Distribution of cardiovascular related cells within the human thalamus. Clinical Autonomic Research, 8(3), 173–179. 10.1007/BF02281122

Park, H.-D., Barnoud, C., Trang, H., Kannape, O. A., Schaller, K., & Blanke, O. (2020). Breathing is coupled with voluntary action and the cortical readiness potential. Nature Communications, 11(1), 289. 10.1038/s41467-019-13967-9

Park, H.-D., Bernasconi, F., Bello-Ruiz, J., Pfeiffer, C., Salomon, R., & Blanke, O. (2016). Transient Modulations of Neural Responses to Heartbeats Covary with Bodily Self-Consciousness. Journal of Neuroscience, 36(32), 8453–8460. 10.1523/JNEUROSCI.0311-16.2016

Park, H.-D., Bernasconi, F., Salomon, R., Tallon-Baudry, C., Spinelli, L., Seeck, M., Schaller, K., & Blanke, O. (2018). Neural Sources and Underlying Mechanisms of Neural Responses to Heartbeats, and their Role in Bodily Self-consciousness: An Intracranial EEG Study. Cerebral Cortex, 28(7), 2351–2364. 10.1093/cercor/bhx136

Park, H.-D., & Blanke, O. (2019). Coupling Inner and Outer Body for Self-Consciousness. Trends in Cognitive Sciences, 23(5), 377–388. 10.1016/j.tics.2019.02.002

Park, H.-D., Correia, S., Ducorps, A., & Tallon-Baudry, C. (2014). Spontaneous fluctuations in neural responses to heartbeats predict visual detection. Nature Neuroscience, 17(4), 612–618. 10.1038/nn.3671

Pattinson, K., Mitsis, G., Harvey, A., Jbabdi, S., Dirckx, S., Mayhew, S., Rogers, R., Tracey, I., & Wise, R. (2009). Determination of the human brainstem respiratory control network and its cortical connections in vivo using functional and structural imaging. NeuroImage, 44(2), 295–305. 10.1016/j.neuroimage.2008.09.007

Pattinson, K. T. S., Governo, R. J., MacIntosh, B. J., Russell, E. C., Corfield, D. R., Tracey, I., & Wise, R. G. (2009). Opioids Depress Cortical Centers Responsible for the Volitional Control of Respiration. Journal of Neuroscience, 29(25), 8177–8186. 10.1523/JNEUROSCI.1375-09.2009

Paulk, A. C., Kfir, Y., Khanna, A. R., Mustroph, M. L., Trautmann, E. M., Soper, D. J., Stavisky, S. D., Welkenhuysen, M., Dutta, B., Shenoy, K. V., Hochberg, L. R., Richardson, R. M., Williams, Z. M., & Cash, S. S. (2022). Large-scale neural recordings with single neuron resolution using Neuropixels probes in human cortex. Nature Neuroscience, 25(2), 252–263. 10.1038/s41593-021-00997-0

Pazo, J. H., & Belforte, J. E. (2002). Basal ganglia and functions of the autonomic nervous system. Cellular and Molecular Neurobiology, 22(5–6), 645–654. 10.1023/a:1021844605250

Perakakis, P. (2019). HEPLAB: A Matlab graphical interface for the preprocessing of the heartbeat-evoked potential (v1.0.0) [Computer software]. Zenodo. 10.5281/ZENODO.2649943

Perl, O., Ravia, A., Rubinson, M., Eisen, A., Soroka, T., Mor, N., Secundo, L., & Sobel, N. (2019). Human non-olfactory cognition phase-locked with inhalation. Nature Human Behaviour, 3(5), 501–512. 10.1038/s41562-019-0556-z

Pinsky, M. R. (2006). Heart–Lung Interactions. In Clinical Critical Care Medicine (pp. 369– 382). Elsevier. 10.1016/B978-0-323-02844-8.50039-1

Pollatos, O., & Schandry, R. (2004). Accuracy of heartbeat perception is reflected in the amplitude of the heartbeat-evoked brain potential: Heartbeat-evoked potential and heartbeat perception. Psychophysiology, 41(3), 476–482. 10.1111/1469-8986.2004.00170.x

Quian Quiroga, R., Nadasdy, Z., & Ben-Shaul, Y. (2004). Unsupervised Spike Detection and Sorting with Wavelets and Superparamagnetic Clustering. Neural Computation, 16(8), 1661–1687. 10.1162/089976604774201631

Raeva, S. (1986). Localization in human thalamus of units triggered during ‘verbal commands,’ voluntary movements and tremor. Electroencephalography and Clinical Neurophysiology, 63(2), 160–173. 10.1016/0013-4694(86)90009-X

Rassler, B., & Raabe, J. (2003). Co-ordination of breathing with rhythmic head and eye movements and with passive turnings of the body. European Journal of Applied Physiology, 90(1–2), 125–130. 10.1007/s00421-003-0876-5

Ringach, D. L., Bredfeldt, C. E., Shapley, R. M., & Hawken, M. J. (2002). Suppression of Neural Responses to Nonoptimal Stimuli Correlates With Tuning Selectivity in Macaque V1. Journal of Neurophysiology, 87(2), 1018–1027. 10.1152/jn.00614.2001

Rinvik, E., & Ottersen, O. P. (1993). Terminals of subthalamonigral fibres are enriched with glutamate-like immunoreactivity: An electron microscopic, immunogold analysis in the cat. Journal of Chemical Neuroanatomy, 6(1), 19–30. 10.1016/0891-0618(93)90004-N

Salehi, S., A. Dehaqani, M. R., Noudoost, B., & Esteky, H. (2020). Distinct mechanisms of face representation by enhancive and suppressive neurons of the inferior temporal cortex. Journal of Neurophysiology, 124(4), 1216–1228. 10.1152/jn.00203.2020

Saper, C. B. (2002). The Central Autonomic Nervous System: Conscious Visceral Perception and Autonomic Pattern Generation. Annual Review of Neuroscience, 25(1), 433–469. 10.1146/annurev.neuro.25.032502.111311

Schäfer, A., & Kratky, K. W. (2008). Estimation of Breathing Rate from Respiratory Sinus Arrhythmia: Comparison of Various Methods. Annals of Biomedical Engineering, 36(3), 476–485. 10.1007/s10439-007-9428-1

Schandry, R., & Montoya, P. (1996). Event-related brain potentials and the processing of cardiac activity. Biological Psychology, 42(1–2), 75–85. 10.1016/0301-0511(95)05147-3

Schandry, R., Sparrer, B., & Weitkunat, R. (1986). From the heart to the brain: A study of heartbeat contingent scalp potentials. International Journal of Neuroscience, 30(4), 261–275. 10.3109/00207458608985677

Schulz, A., Schilling, T. M., Vögele, C., Larra, M. F., & Schächinger, H. (2016). Respiratory modulation of startle eye blink: A new approach to assess afferent signals from the respiratory system. Philosophical Transactions of the Royal Society B: Biological Sciences, 371(1708), 20160019. 10.1098/rstb.2016.0019

Seth, A. K. (2013). Interoceptive inference, emotion, and the embodied self. Trends in Cognitive Sciences, 17(11), 565–573. 10.1016/j.tics.2013.09.007

Seyal, M., & Bateman, L. M. (2009). Ictal apnea linked to contralateral spread of temporal lobe seizures: Intracranial EEG recordings in refractory temporal lobe epilepsy. Epilepsia, 50(12), 2557–2562. 10.1111/j.1528-1167.2009.02245.x

Shao, S., Shen, K., Wilder-Smith, E. P. V., & Li, X. (2011). Effect of pain perception on the heartbeat evoked potential. Clinical Neurophysiology: Official Journal of the International Federation of Clinical Neurophysiology, 122(9), 1838–1845. 10.1016/j.clinph.2011.02.014

Solcà, M., Park, H.-D., Bernasconi, F., & Blanke, O. (2020). Behavioral and neurophysiological evidence for altered interoceptive bodily processing in chronic pain. NeuroImage, 217, 116902. 10.1016/j.neuroimage.2020.116902

Sverrisdóttir, Y. B., Green, A. L., Aziz, T. Z., Bahuri, N. F. A., Hyam, J., Basnayake, S. D., & Paterson, D. J. (2014). Differentiated Baroreflex Modulation of Sympathetic Nerve Activity During Deep Brain Stimulation in Humans. Hypertension, 63(5), 1000–1010. 10.1161/HYPERTENSIONAHA.113.02970

Tadel, F., Baillet, S., Mosher, J. C., Pantazis, D., & Leahy, R. M. (2011). Brainstorm: A User-Friendly Application for MEG/EEG Analysis. Computational Intelligence and Neuroscience, 2011, 1–13. 10.1155/2011/879716

Takada, M., Tokuno, H., Hamada, I., Inase, M., Ito, Y., Imanishi, M., Hasegawa, N., Akazawa, T., Hatanaka, N., & Nambu, A. (2001). Organization of inputs from cingulate motor areas to basal ganglia in macaque monkey: Cingulate motor areas inputs to basal ganglia. European Journal of Neuroscience, 14(10), 1633–1650. 10.1046/j.0953-816x.2001.01789.x

Task Force, T. F. of T. E. S. of C. and T. N. A. S. of P. and E. (1996). Heart rate variability: Standards of measurement, physiological interpretation and clinical use. Task Force of the European Society of Cardiology and the North American Society of Pacing and Electrophysiology. Circulation, 93(5), 1043–1065.

Terem, I., Dang, L., Champagne, A., Abderezaei, J., Pionteck, A., Almadan, Z., Lydon, A., Kurt, M., Scadeng, M., & Holdsworth, S. J. (2021). 3D amplified MRI (aMRI). Magnetic Resonance in Medicine, 86(3), 1674–1686. 10.1002/mrm.28797

Thompson, R. O. (1979). Coherence significance levels. J. Atmos. Sci., 36, 2020–2021.

Thornton, J. M., Aziz, T., Schlugman, D., & Paterson, D. J. (2002). Electrical stimulation of the midbrain increases heart rate and arterial blood pressure in awake humans. The Journal of Physiology, 539(2), 615–621. 10.1113/jphysiol.2001.014621

Tsakiris, M., & Critchley, H. (2016). Interoception beyond homeostasis: Affect, cognition and mental health. Philosophical Transactions of the Royal Society B: Biological Sciences, 371(1708), 20160002. 10.1098/rstb.2016.0002

Usrey, W. M., & Alitto, H. J. (2015). Visual Functions of the Thalamus. Annual Review of Vision Science, 1(1), 351–371. 10.1146/annurev-vision-082114-035920

Vibert, J. F., Caille, D., Bertrand, F., Gromysz, H., & Hugelin, A. (1979). Ascending projection from the respiratory centre to mesencephalon and diencephalon. Neuroscience Letters, 11(1), 29–33. 10.1016/0304-3940(79)90051-X

Weiss, N., Ohara, S., Johnson, K. O., & Lenz, F. A. (2009). The Human Thalamic Somatic Sensory Nucleus [Ventral Caudal (Vc)] Shows Neuronal Mechanoreceptor-Like Responses to Optimal Stimuli for Peripheral Mechanoreceptors. Journal of Neurophysiology, 101(2), 1033–1042. 10.1152/jn.90990.2008

Zavala, B., Damera, S., Dong, J. W., Lungu, C., Brown, P., & Zaghloul, K. A. (2015). Human Subthalamic Nucleus Theta and Beta Oscillations Entrain Neuronal Firing During Sensorimotor Conflict. Cerebral Cortex, bhv244. 10.1093/cercor/bhv244

Zelano, C., Jiang, H., Zhou, G., Arora, N., Schuele, S., Rosenow, J., & Gottfried, J. A. (2016). Nasal Respiration Entrains Human Limbic Oscillations and Modulates Cognitive Function. The Journal of Neuroscience, 36(49), 12448–12467. 10.1523/JNEUROSCI.2586-16.2016

