## Supplemental_information for "Single neurons in thalamus and subthalamic nucleus process cardiac and respiratory signals in humans"

### Supplementary material

#### Identification of EAPs modulated by cardioballistic effect

A big challenge in evaluating neural activity relative to the cardiac cycle is the presence of heart cycle-related artifacts in electrophysiological recordings. The presence of artifacts, such as the cardiac field artifact (CFA) or the pulse artifact (PA), has been extensively investigated for scalp EEG data, but less-so for intracranial recordings (Kern et al., 2013). Intraoperative electrophysiological monitoring is known to be affected by the cardioballistic effect, observed in the waxing and waning of the extracellular recorded signal (Montgomery, Jr, 2014). This effect is associated with pulse waves that reach blood vessels in the target region and cyclically move the tip of the electrode relative to the recorded neuron. The cardioballistic effect seems to be more prominent in subcortical regions, possibly because of the presence of extended vascularization and is reduced following propofol administration that reduces the patient's blood pressure (MacIver et al., 2011). Recent work (Mosher et al., 2020) has highlighted how cardiac cycle artifacts are present in different brain regions and can affect spike identification by modulating the extracellular action potential (EAP) detected and can create spurious modulations of neural activity. We applied their methodology to identify single units in our recordings that presented an EAP modulation. For each neuron identified, we calculated the average EAP waveform along the cardiac cycle and evaluated how 4 different EAPs features changed along the cycle. Neurons that presented at least one significant cyclically modulated EAP feature (according to a statistical surrogate test, see Materials and Methods) were labeled as artifacted and excluded from further analysis. Following this analysis, 42% of units (54 out of 127) presented a significant EAP modulation and were therefore excluded from further analysis. Examples of a modulated and unmodulated neuron are shown in supplementary Figure S1. For each tested neuron we also extracted the Motion Index (MI) z-scored to the surrogate distribution. We then tested the distribution of z-scored MIs against zero separately for EAP non-modulated (stable) and EAP modulated (likely affected by artifacts) units. We found that the distributions on each of the four EAP features were not different from zero for the subpopulation of EAP non-modulated neurons, while they were significantly larger than zero for the population of EAP modulated units (t-test Stats: AMP non-modulated: $t(72) = 0.36$, $p = 0.7$; HW non-modulated: $t(72) = 1.67$, $p = 0.1$; TPW non-modulated $t(72) = 1.08$, $p = 0.28$; REP non-modulated: $t(72) = 0.5$, $p = 0.61$; AMP modulated: $t(53) = 11.9$, $p = 1.2e-16$; HW modulated: $t(53) = 6.83$, $p = 8.2e-9$ ; TPW modulated $t(53) = 4.26$, $p = 8.4e-5$; REP modulated: $t(53) = 6.18$, $p = 9.4E-8$). These results confirm that the EAP waveform modulation test was able to identify a sub-population of units (the ones included in the analysis), for which the motion indices were not different from zero.

### Figure S1

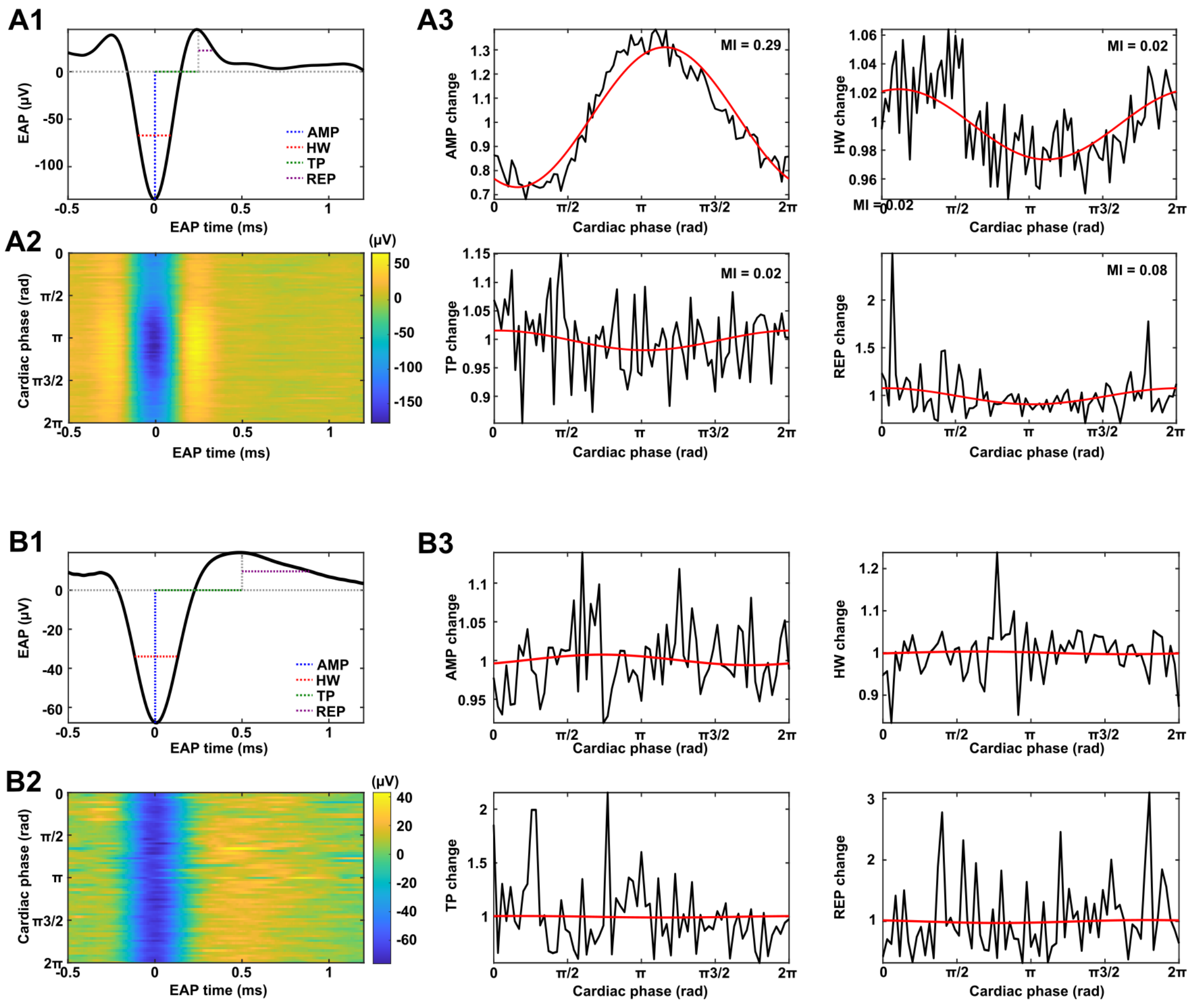


**Figure S1. EAP modulation test. A)** Example of a neuron with ECG modulated EAPs recorded from Vc. **A1)** Average EAP with highlighted (color-coded) the four features considered for the motion test. **A2)** Color-plot of EAP shape along the cardiac cycle (in rad). **A3)** Each panel shows the variation of a specific feature (AMP, HW, TP, REP) along the cardiac cycle relative to the mean value of the feature. The red curve shows the cosine fit of the data used to calculate the motion index. **B)** Example of a neuron with stable EAPs recorded in STN. Conventions as in panels A. Note that even though the selected example neurons were recorded from different target locations, the proportion of EAP modulated neurons was similar across regions and there is no statistical difference across the populations of modulation indices measured in each region for any of the 4 features (one-way ANOVA: AMP $F(2,124) = 0.5$, $p = 0.6$; HW $F(2,124) = 0.9$, $p = 0.41$; TPW $F(2,124) = 2.1$, $p = 0.13$; REP $F(2,124) = 2.4$, $p = 0.09$).

### Figure S2

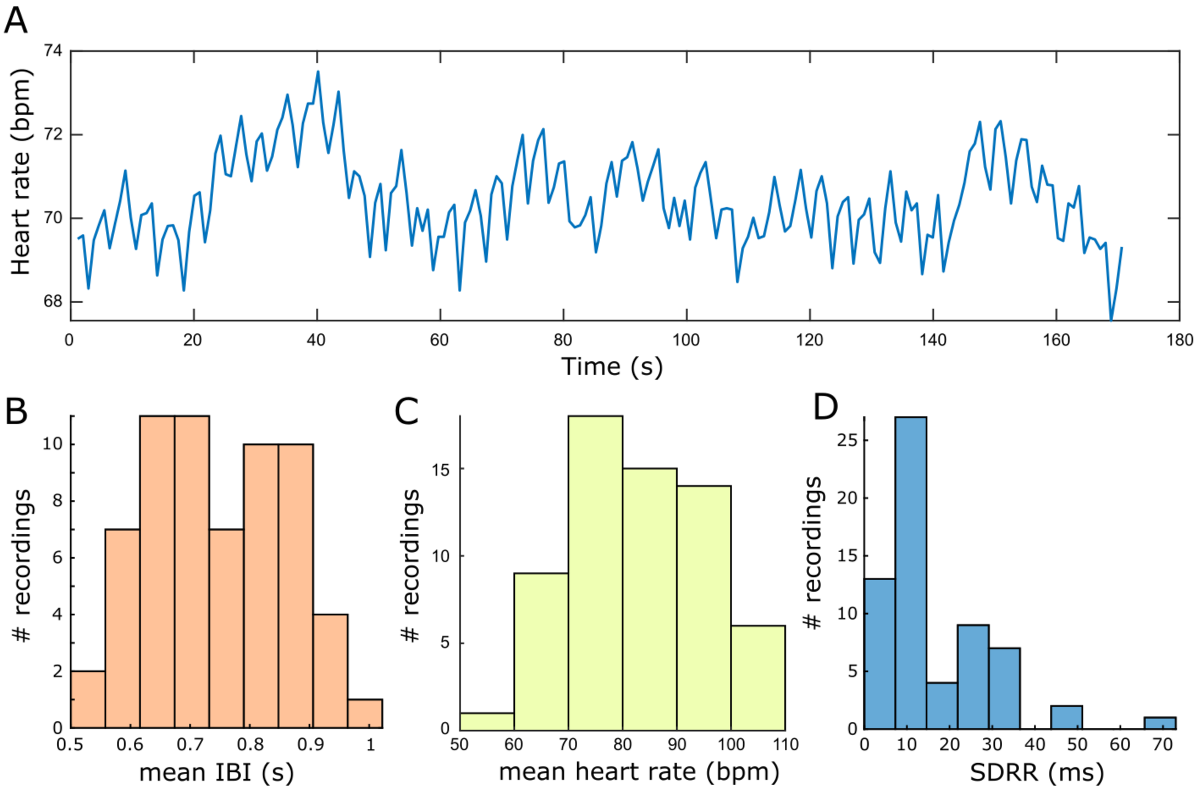


**Figure S2. Heart rate variability across the 63 recordings. A)** Exemplary heart rate (HR) trace acquired during one recording session **B)** Distribution of mean IBIs (average value across recordings = 0.75s (STD =0.1s). **C)** Distribution of average HRs in bpm across recording sessions (average value across recordings = 82.3bpm, SD = 12.8bpm). **D)** Distribution of SDRR (standard deviation of the IBIs for all beats).

### Figure S3

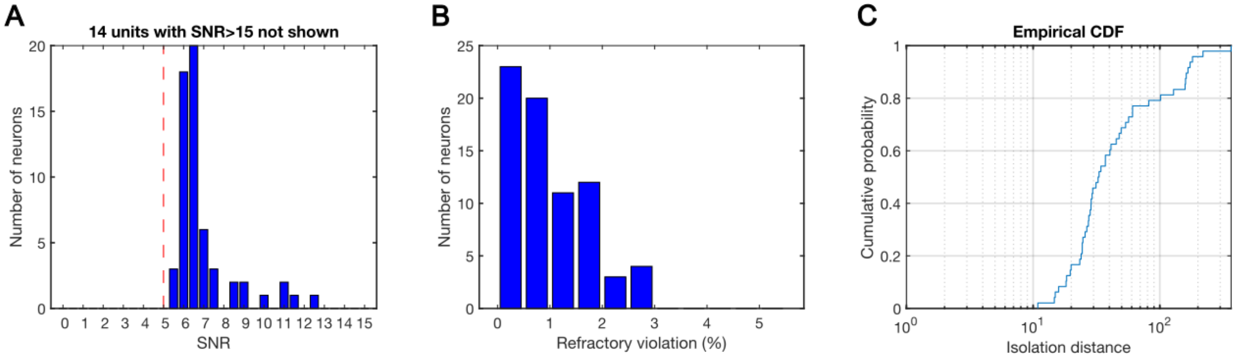


**Figure S3. Spikes quality metrics across the 73 single units included in the analysis. A)** Distribution of spikes' signal-to-noise ratio (SNR). Red dotted line indicates the detection threshold used by our detection algorithm. 14 neurons with SNR > 15 are not shown in the plot to ease the visualization of the distribution. **B)** Percentage of refractory violation (percentage of Inter-Spike intervals falling within the 3ms interval). **C)** Empirical cumulative distribution of Isolation distances for identified single neurons.

### Figure S4.1

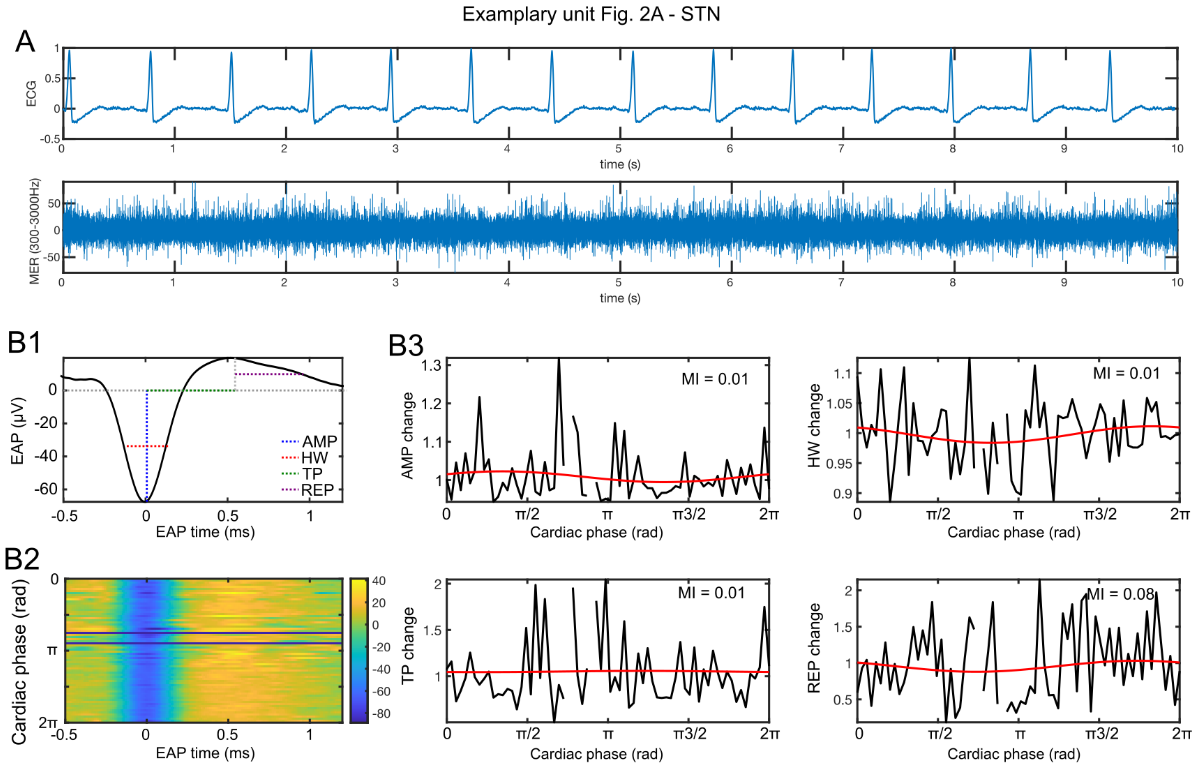


**Figure S4.1. Sample recordings and EAP waveform modulation test for Exemplary unit in Figure 2 (left) recorded from the STN**. **A)** Sample ECG (top) and raw high-frequency MER. **B1)** Average EAP with highlighted (color-coded) the four features considered for the motion test. **B2)** Color-plot of EAP shape along the cardiac cycle (in rad). **B3)** Each panel shows the variation of a specific feature (AMP, HW, TP, REP) along the cardiac cycle relative to the mean value of the feature. The red curve shows the cosine fit of the data used to calculate the motion index. Motion indices z-scored to the surrogate population are as follows: MI-AMP = -2.1, MI-HW = 0.059, MI-TP = -1.9, MI-REP = 2.3.

### Figure S4.2

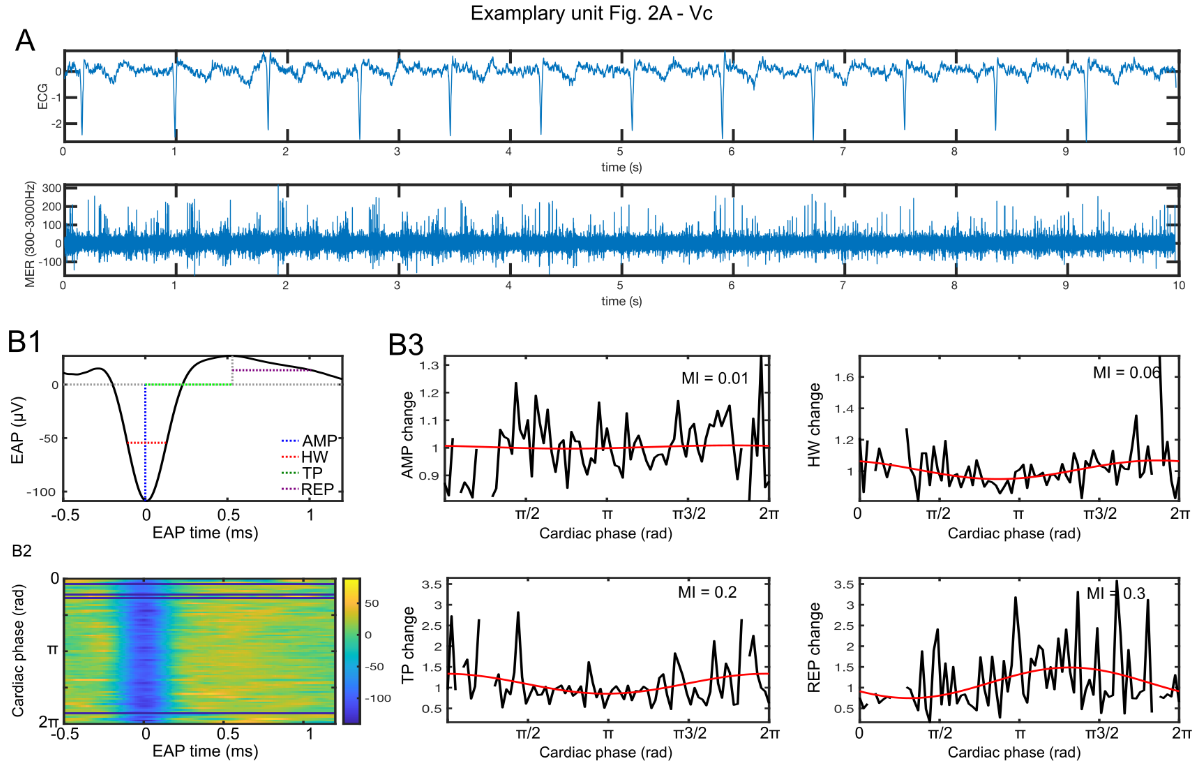


**Figure S4.2. Sample recordings and EAP waveform modulation test for Exemplary unit in Figure 2 (center) recorded from Vc.** Conventions are as in Figure S4.1. Motion indices z-scored to the surrogate population are as follows: MI-AMP = -0.51, MI-HW = 0.67, MI-TP = -0.33, MI-REP = 1.8.

### Figure S4.3

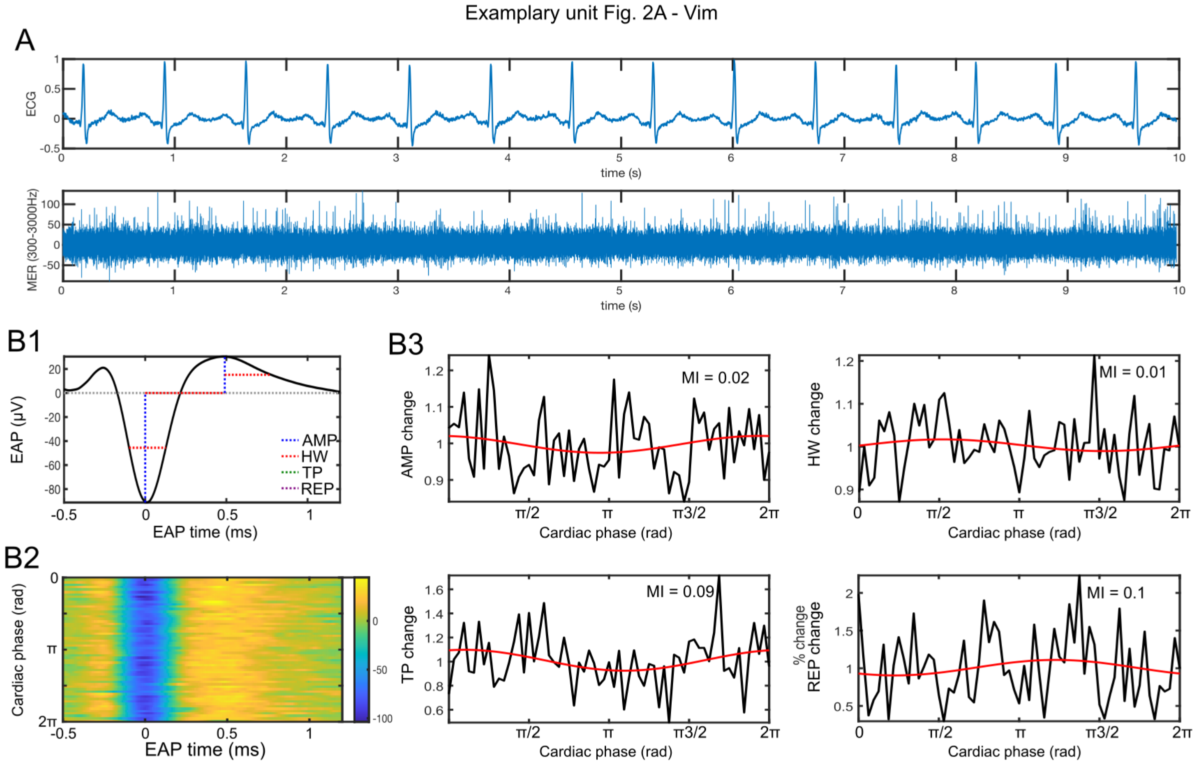


**Figure S4.3. Sample recordings and EAP waveform modulation test for Exemplary unit in Figure 2 (right) recorded from Vim.** Conventions are as in Figure S4.1. Motion indices z-scored to the surrogate population are as follows: MI-AMP = 1.5, MI-HW = 1.3, MI-TP = -0.69, MI-REP = 2.1.

#### Signal-to-noise ratio

Signal-to-noise ratio (SNR) is an important measure to determine whether the heartbeat responses observed here could be explained by spikes near the detection threshold. To control for this possibility, we compared SNR between heartbeat responsive and non-responsive units. For each of the units included in the analysis we calculated the SNR as the mean amplitude of the waveform over the background noise. Background noise ($\sigma_N$) was estimated from the filtered signal as done by the detection algorithm we used (wave_clus get_spikes). We found that all units were well above the detection threshold used ($5\sigma_N$) and that there was no difference in terms of SNR between units that were responsive and non-responsive to the heartbeat (Figure S5, Wilcoxon rank sum, $Z = 0.88$, $p = 0.38$; note that we used non-parametric statistics as the distributions were clearly skewed). This result indicates that responsiveness to the heartbeat cannot be explained by misdetection of spikes near the threshold; because if that would have been the case, we would have expected to find that responsive units had significantly lower SNR than non-responsive ones.

### Figure S5

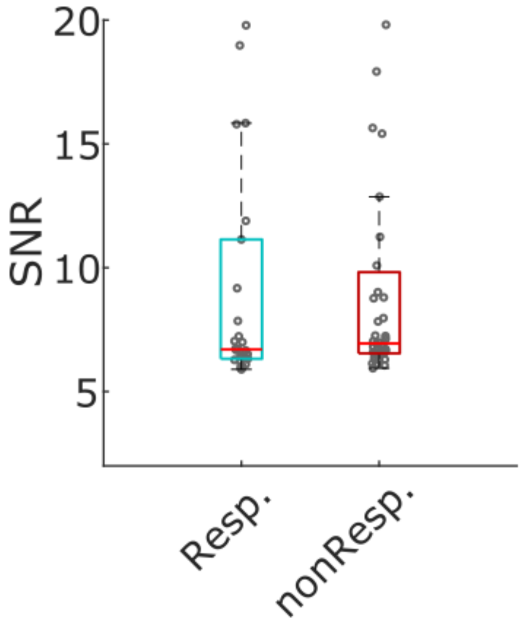


**Figure S5**. **Signal-to-noise ratio and heartbeat responses**. Boxplot of signal-to-noise ratio (SNR) for each neuron included in the analysis separated in heartbeat responsive and non-responsive. Horizontal line shows median, box indicates the 25th and 75th percentiles, respectively. No difference in SNR were observed between the two groups (Wilcoxon rank sum, Z = 0.88, p = 0.38)

#### Laterality analyses

After the identification of neurons responsive to the heartbeat, we further verified whether aspects of the neural responses to the heartbeat could differ between recording regions in the left vs the right hemisphere. We note that our dataset included more neurons recorded from the left side (48 units on the left vs 25 on the right). However, there was no significant difference in the proportion of neurons responsive to the heartbeat between the two sides according to a z-test on proportions (22 units on the left and 8 on the right, $p$=0.25). Furthermore, we observed no difference in the amplitude of responses between units recorded from the left vs right side (unpaired t-test, $t(28) = 1.1$, $p = 0.28$). The amplitude of response for each heartbeat responsive unit was computed as the maximal change of firing rates along the cardiac cycle divided by the mean firing rate of the neuron. Finally, when we analyzed the peak latency of such modulation for the neurons responsive to the heartbeat, we observed no significant difference between peak latencies on the left and right side (unpaired t-test, t(28) $=1.3$, $p = 0.21$).

#### Lagged correlation

We asked whether the identified relationship between firing rate and IBI (Figure 3) extended to several cardiac cycles. We looked for longer lasting effects of correlation that could span across several cardiac cycles by analyzing the lagged correlation between the average firing rate during each cardiac cycle (FRc) and IBIs when IBIs values were shifted by a lag between -10 and 10 cycles. Once we identified the lag with maximum cross-correlation between the two measures (max lag), we then calculated Pearson's correlation between FRc and the IBI shifted by the max lag value to assess significance (FDR corrected for multiple comparisons). We identified only 8 neurons (out of the 73, corresponding to ~11%) for which the maximal significant correlation happened at a lag different from zero. Specifically, for 4 units we found a positive lag, indicating that changes in the firing rate precede changes in the IBI, while for 4 units we found a negative lag, suggesting the opposite direction of change. The number of neurons with positive vs negative Max Lag across regions is shown in Figure S6.

### Figure S6

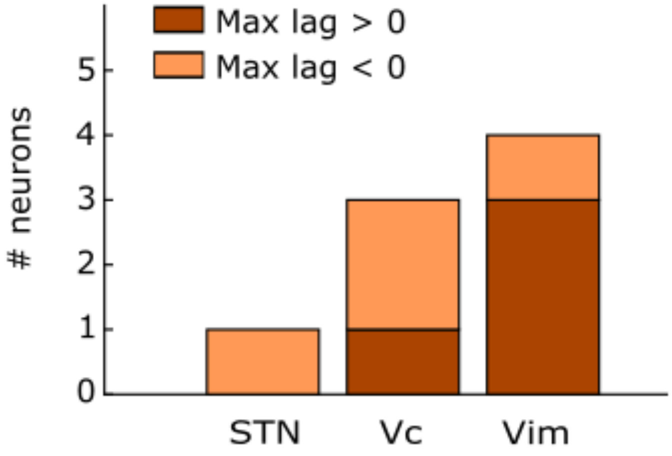


**Figure S6. Lagged correlation neurons.** Distribution of neurons exhibiting a significant lagged correlation between IBI and firing rate in the three regions (with a lag different from 0, dark orange for positive lags, light orange for negative lags).

### Figure S7

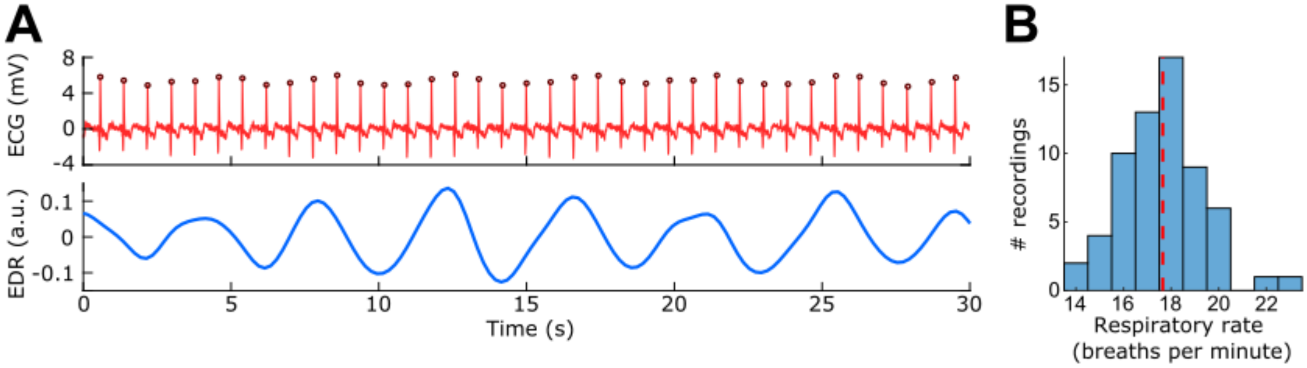


**Figure S7. Example of ECG-Derived Respiration (EDR) and average respiratory rates across recording sessions. A)** Top panel shows a 30 seconds ECG signal with a clear modulation of the R-peak amplitude due to the cyclic changes in impedance between the heart and ECG electrodes caused by the respiration. Circles overlaid on ECG signal indicate R-peaks. Bottom panel shows the respiratory signal derived from the ECG in the same time interval using the feature-based EDR algorithm. **B)** - Distribution of Respiratory Rates measures across recording sessions (mean =17.7 breaths per minute, SD = 1.7).

### Multimodal responses and RSA

Respiratory sinus arrythmia (RSA) reflects coupling between respiratory and cardiac signals. It is in principle possible that individuals with increased RSA may exhibit increased cardio-respiratory coupling at the level of single neurons too. To test this hypothesis, we measured RSA in our dataset (distribution shown below Figure S8-A) and then split the sessions in two groups of low vs high RSA based on the median value (1.73bpm). RSA was estimated using a peak-valley procedure (Grossman et al., 1990) and is expressed in beats per minute (bpm). We then verified whether the proportion of multimodal (cardiac + respiratory) units were different in the low vs high RSA groups. Among the 13 multimodal units observed, 7 were recorded from low RSA sessions and 6 were recorded from high RSA sessions. Furthermore, the proportion of unimodal (cardiac only or respiratory only) vs multimodal (cardiac and respiratory) units were comparable in both low and high RSA groups (shown below, Figure S8-B) [Low RSA: 19/7 (unimodal/multimodal); High RSA: 21/6 (unimodal/multimodal). These results strongly suggest that the multimodal neurons observed here are not likely to respond to the coupling between cardiac and respiratory signals.

### Figure S8

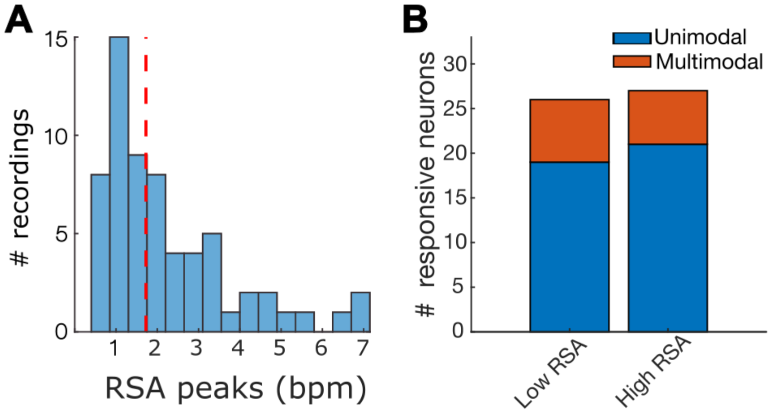


**Figure S8. Multimodal responses and RSA. A)** Distribution of respiratory sinus arrhythmia (RSA) measured across the recording sessions (red dotted line indicates median of the distribution =1.73 bpm). **B)** Numbers of unimodal and multimodal responses detected in session with low and high RSA.

### Table S1

| Patient | # recordings | # neurons (after artifact removal) | | |
|---|---|---|---|---|
| | | STN | Vc | Vim |
| **1** | 4 | 0 | 4 | 2 |
| **2** | 3 | 0 | 1 | 0 |
| **3** | 1 | 1 | 0 | 0 |
| **4** | 2 | 0 | 0 | 5 |
| **5** | 2 | 0 | 0 | 0 |
| **6** | 4 | 0 | 3 | 1 |
| **7** | 6 | 6 | 0 | 0 |
| **8** | 2 | 0 | 0 | 3 |
| **9** | 2 | 1 | 0 | 0 |
| **10** | 9 | 0 | 8 | 3 |
| **11** | 2 | 0 | 2 | 0 |
| **12** | 4 | 0 | 4 | 1 |
| **13** | 2 | 2 | 0 | 0 |
| **14** | 2 | 0 | 0 | 3 |
| **15** | 2 | 2 | 0 | 0 |
| **16** | 2 | 0 | 0 | 5 |
| **17** | 2 | 0 | 2 | 0 |
| **18** | 2 | 1 | 0 | 0 |
| **19** | 2 | 0 | 0 | 6 |
| **20** | 2 | 0 | 0 | 3 |
| **21** | 1 | 0 | 0 | 1 |
| **22** | 1 | 1 | 0 | 0 |
| **23** | 4 | 2 | 0 | 0 |

Table S1. Number of recording chunks and neurons in each area per patient.

### References

Grossman, P., Beek, J., Wientjes, C., 1990. A Comparison of Three Quantification Methods for Estimation of Respiratory Sinus Arrhythmia. Psychophysiology 27, 702–714. https://doi.org/10.1111/j.1469-8986.1990.tb03198.x

Kern, M., Aertsen, A., Schulze-Bonhage, A., Ball, T., 2013. Heart cycle-related effects on event-related potentials, spectral power changes, and connectivity patterns in the human ECoG. NeuroImage 81, 178–190. https://doi.org/10.1016/j.neuroimage.2013.05.042

MacIver, M.B., Bronte-Stewart, H.M., Henderson, J.M., Jaffe, R.A., Brock-Utne, J.G., 2011. Human Subthalamic Neuron Spiking Exhibits Subtle Responses to Sedatives. Anesthesiology 115, 254–264. https://doi.org/10.1097/ALN.0b013e3182217126

Montgomery, Jr, E.B., 2014. Intraoperative Neurophysiological Monitoring for Deep Brain Stimulation: Principles, Practice and Cases. Oxford University Press. https://doi.org/10.1093/med/9780199351008.001.0001

Mosher, C.P., Wei, Y., Kamiński, J., Nandi, A., Mamelak, A.N., Anastassiou, C.A., Rutishauser, U., 2020. Cellular Classes in the Human Brain Revealed In Vivo by Heartbeat-Related Modulation of the Extracellular Action Potential Waveform. Cell Rep. 30, 3536-3551.e6. https://doi.org/10.1016/j.celrep.2020.02.027
